# Host signature as major driver of root and rhizosphere core microbiomes that differently affect plant functional traits

**DOI:** 10.1101/2023.11.23.568511

**Authors:** Jipeng Luo, Yuanfan Wang, Yu Zhang, Wenzhe Gao, Yongchao Liang, He-Ping Zhao, Shaohua Gu, Tingqiang Li

## Abstract

**Background:** Plant can evolve with a core root microbiome that maintains essential functions for host performance. However, the relative importance of plant traits and soil factors on the structure, assembly, co-occurrence networks of the core root microbiomes and their relevance for plant characteristics remain elusive. Here, we investigated how plant species identity and soil environment affect the core bacterial communities in the bulk soil, rhizosphere and root endosphere of four plants with a gradient of Cd/Zn accumulation capacity under controlled and field environments. We further tested on the role of the core bacterial isolates in plant growth and accumulation of metal and nutrients.

**Results:** We identified root compartment and plant species rather than environmental parameters as the primary driver of Cd-accumulator root microbiome. Stochastic processes were more important for the assembly of endosphere generalists (58.5%) than rhizosphere counterparts (45.2%), indicating that generalists were more robust to environmental changes. Increasing host selection from epiphytes to endophytes resulted in the existence of the endosphere and rhizosphere generalist core microbiota common to different plants under varying growth environments, highlighting that shared environmental and physiological features of host plants are decisive for core microbiome establishment. Further, endophytic core microbiota conferred greater biotic connectivity within networks and was more important predictors of plant metal accumulation, whereas the rhizosphere cores were more closely linked to plant biomass and nutrient status. The divergent functions of rhizosphere and endosphere core microbes on plant characteristics were also validated by inoculating the synthetic communities comprising bacterial isolates belonging to the core microbiota.

**Conclusion:** This study indicated the pivotal role of plant trait in the assembly of conserved and functionally important core microbiome common to different Cd-accumulators, which brings us closer to manipulating the persistent root microbial associations to accelerate the rejuvenation of metal-disturbed soils through host genetics.

## Background

The host plant and its associated microbiota form a ‘holobiont’, whereby they have been deeply intertwining and co-evolving, contributing to the overall stability of the system [1]. Root-associated microbiota can be considered an extension of the host’s genome and is known to promote host plant productivity and fitness, modulate developmental pathways and confer plant tolerance to abiotic stresses [2–4]. Empirical evidence suggests that the phenotypic expression of host traits is due to the combined genetic expression of both the host and the host-associated microbiome [5, 6]. These studies have greatly increased interest in the potential to harness root microbiome to enhance plant phenotypes involved in plant resilience and yield [7, 8]. However, a roadblock hampering this effort is a lack of understanding of which microbes can consistently associate with the plants across different environments, the drivers of the persistent microbial community, and the plant-supportive functions of the persistent microbiomes inhabiting distinct root habitats.

Plant root microbiomes are acquired horizontally from the soil reservoir, for which memberships are largely driven by environmental factors such as soil type, geography, land use history and pollutants [9–12]. The host plant exerts additional selective pressure on its microbiota, resulting in filtered subsets of soil microbes often composed of consistently enriched microbial taxa on and inside roots [13]. Given that microbiota can have multiple impacts on plant growth and functional characteristics [14, 15], particularly under different environmental conditions [16, 17], it is likely that the filtering process is under selection and may lead to microbe-mediated adaptation and phenotypic expression [15, 16]. Notably, most studies investigating the effect of host species identity on root and rhizosphere microbiota have focused exclusively on different genotypes, ecotypes or cultivars within a single plant species. However, little information is available on the degree to which microbial communities living in association with phylogenetically divergent hosts overlap with each other.

Although many members of the plant microbiota are commensals, a small subset provides beneficial services for plant phenotypic traits and fitness [13, 18]. Recent studies have shown that plant functional traits (morphological and physiological characteristics), such as specific leaf area, root system architecture, below- and above-ground biomass, can be engineered by the composition of the root microbiome and its beneficial interactions with host plants [6, 19, 20]. However, the root microbiome is highly diverse and dynamic, with the majority of its members associating with the plants transiently and opportunistically, and therefore unlikely to contribute significantly to plant’s biological functions [21, 22]. Hence, there is an increasing need to filter out transient microbiota to refine focus on the core microbiota, which consists of persistent microbiome members that are associated with the plant and thus have a greater likelihood of affecting the host phenotype [21–23]. Generalist species can be defined as taxa that are able to adapt to a wide range of habitats and environments [24], and have a general characteristic of great metabolic flexibility that allows them to compete in highly dynamic and even suboptimal environments [25]. Thus, the persistent and stable distribution of generalist species enables them to be ideal candidates of the core microbiome, i.e. taxa with high occupancy that persist across multiple microbial communities associated with a habitat [19, 26]. Although the importance of core microbiomes in plant fitness and growth has been previously investigated [27, 28], their ecological role in the stability of rhizosphere and endosphere co-occurrence networks and plant functional traits remains poorly understood.

The increasing contamination of agricultural soils with heavy metal(loid)s due to anthropogenic activities poses huge threats to the diversity and metabolic potential of soil microbiomes and associated ecosystem functions, ranging from biogeochemical cycle to the maintenance of plant growth [29, 30]. Heavy metals, such as cadmium (Cd) and zinc (Zn), are two of the most ubiquitous and bioavailable heavy metals in rice paddy soils, leading to concerns that the rice produced is metal contaminated [31, 32]. Phytoextraction, the use of metal (hyper)accumulating plants to remove metal(loid)s from soils by concentrating them in the harvestable parts, has been proven as an eco-friendly and cost-effective alternative to remediate contaminated soils [33] and has received global attention due to its potential to restore soil quality and some functional capacities [34, 35]. Moreover, a growing number of studies have shown that the association of hyperaccumulators with different microbial communities results in contrasting phytoremediation ability [36–38]. As a result, the metal hyperaccumulating plant has emerged as a model for disentangling the links between core microbiomes and plant functional characteristics, such as plant uptake and accumulation of heavy metals. Therefore, insights into the structure of generalist core microbiomes between Cd-accumulators and the factors that affect them, as well as the role of these core cohorts in plant growth, metal and nutrient accumulation, would lay a theoretical foundation for developing rational microbiome-based strategies for the enhancing the efficacy of phytoextraction via host genetics.

To achieve our goals, we first analyzed bacterial community composition in root endosphere, rhizosphere and bulk soil associated with four different Cd-accumulators with a varying Cd/Zn accumulation ability, grown in the three types of paddy soils with variable Cd concentrations and distinct geographical origins under field and controlled environments. Then, the ecological assembly processes, co-occurrence networks and potential functions of habitat generalists, specialists and generalist core microbiota along the soil-root continuum were investigated. Finally, to assess the capability of core microbial populations to promote plant growth and facilitate metal and nutrient accumulation, *S. alfredii* plants were inoculated with the SynComs comprising the core rhizospheric or endophytic bacteria isolated from different Cd-accumulators in this study. The current study aimed to address the following questions: (i) How dose the effect of plant species identity compare with that of the environment when structuring the root-associated bacterial microbiota? (ii) Which bacteria are persistent members of the core root-associated microbiota between divergent Cd-accumulators across varying environmental conditions and what are the drivers of such taxa in the core microbiota? (iii) How do the core microbiome members inhabiting the root endosphere and rhizosphere relate to plant functional traits?

## Materials and methods

### Soil collection and characterization

In this study, three types of paddy field soils with varying Cd/Zn pollution levels [highly polluted (HP) vs slightly polluted (SP)] and different geographical origins [HP and SP soils vs non-polluted soil (NP)] were used. Detailed information on the geographical origin, metal pollution level and physicochemical parameters were previously reported in our studies [39, 40] and can also be found in Additional files (Table S1).

### Experimental design, plant growth and sample collection

We investigated the bacterial communities inhabiting the root endosphere, rhizosphere and bulk soil of two Cd-hyperaccumulators, *Sedum alfredii* (*Crassulaceae*) [41] and *Solanum nigrum* (Solanaceae) [42], a Cd accumulator, *Brassica juncea* L. Czern (Brassicaceae) [43], and the non-hyperaccumulating ecotype *S. alfredii* (Crassulaceae) [44] grown in the three paddy soils using a greenhouse experiment. These plants represent a gradient of plant’s ability to extract and remove Cd from the soils in the following order: *S. alfredii* > *S. nigrum* > *B. juncea* > non-hyperaccumulator *S. alfredii*. After soil preparation, plant seeds were surface sterilized with 75% ethanol for 30 s, then in 2.5% sodium hypochlorite three times for 5 min, and germinated in sterile potting medium for three weeks. Twelve treatments were set up across three soils and unplanted pots served as the control treatment, resulting in a total of fifteen treatments. Each treatment was replicated five times, leading to a total of 75 pots arranged in a completely randomised block design. The plants were grown in the greenhouse under natural daylight, with an average day/night temperature of 26/20℃ and relative humidity of 70/80%. The pots were watered regularly every two days to maintain soil moisture at 55-60% of the water holding capacity of each paddy soil. After four months of plant growth, pots were sampled, and shoot, root and rhizosphere and unplanted soil samples were collected as previously described (Additional files, [45]).

### Field experiment

To investigate the impact of the plant growing environment on the core root microbiome, we grew hyperaccumulator *S. alfredii* on two sites from where HP and SP soils were collected. Each site was composed of one block measuring 125 m^2^, and each block contained five plots. *S. alfredii* was selected for this study because of its exceptional capability to extract Cd and Zn from soils compared to other plants and its remarkable root microbial enrichment effect [40], which may represent the core microbiome assembly pattern of other plants. The plants were grown in the paddy field, which began at the time as it grew in the greenhouse, and field management followed local practices. After four months of in situ growth, plants were in good condition. Ten healthy plants of similar size were randomly excavated from each plot and pooled into one sample. Sampling of the root endosphere, rhizosphere and bulk soil and determination of plant metal accumulation, soil metal content and plant nitrogen and phosphorus status were conducted as previously described (Additional files, [39, 40, 45])

### Effects of rhizospheric and endophytic SynComs on plant functional traits

*S. alfredii* plants were grown in clay-based SynComs containing essential nutrient elements for plant growth, inoculating with core bacteria or heat killed core bacteria. The isolation of the bacterial strains belonging to the core rhizosphere and endosphere microbiota was conducted similar to our previously described method [40, 46]. Before plant cultivation, calcined clay, an inert soil substitute, was artificially polluted with Cd (as a solution of CdSO_4_) at a level of 1 mg kg^-1^; such a Cd concentration has been reported to be widely distributed in the paddy field in China [31]. The Cd-spiked soils were aged for 3 months at 60% of water-holding capacity. After the aging period, calcined clay was sterilized three times by autoclaving. Preparation and inoculation of SynComs in calcined clay were similar to a previous study [3]. Briefly, rhizosphere and endosphere bacterial isolates of the core microbiota were cultivated in 50 ml tubes in TSB medium at 28°C for 3 d and subsequently pooled (in equal concentration) to prepare SynComs for inoculations. To inoculate SynComs into the calcined clay, the optical density OD_600_ of SynComs were adjusted to 0.5-1, and 2.0 ml SynComs were mixed in 250 ml 1 × *S. alfredii* growth nutrient solution (pH 5.8) [47], and mixed with 200 g of calcined clay in aseptic plastic boxes (∼10^6^ bacterial cells per g of calcined clay). Then three-week-old sterile *S. alfredii* seedlings were transplanted to the sterile clay subjected to four different treatments, including control (no added SynCom and Cd), Cd treatment, heat killed SynCom plus 1 mg kg^-1^Cd [SynCom(HK)], SynCom plus 1 mg kg^-1^ Cd (SynCom). Each treatment had five independent replicates and each pot contains three seedlings. Plants were harvested after 60 days of growth under a gnotobiotic conditions with a 16 h light cycle, 26/22 °C average day/night temperature and 25% humidity. At harvest, plant root, shoot and soil samples were collected, and shoot fresh weight, root total length, root surface area, plant metal accumulation, plant nitrogen and phosphorus status, and relative expression of transporter genes were measured to evaluate plant growth promotion and microbe-mediated changes in plant functional traits (Additional files, [39, 45, 46]).

### DNA extraction, amplification preparation and bioinformatic analyses

Genomic DNA was extracted from the root endosphere, rhizosphere and bulk soil using the DNeasy® PowerSoil DNA Isolation kit (QIAGEN, Hilden, Germany). The concentration and quality of the extracted DNA were checked using a spectrophotometer (NanoDrop, 2000; Thermo Scientific, USA). Bacterial V3-V4 regions were amplified using a dual-indexed 16S rRNA Illumina iTags primer set of 338F (5′-ACTCCTACGGGAGGCAGCA-3) and 806R (5′-GGACTACHVGGGTWTCTAAT-3) as previously described [48]. A unique 12 bp error-correcting barcode was used to tag each PCR product. Amplicons from all samples were pooled in equimolar ratios and were sequenced on the Illumina HiSeq platform.

A total of 4216175 high-quality sequences were obtained with a median read count per sample of 46846 (range: 24369-60990). Raw reads were processed using the UPARSE pipeline as previously described in our study [39], which included the merging of paired-end reads, quality filtering of de-replicated reads, sequence denoising and taxonomic assignment. Denoising attempts to identify correct biological sequences in the reads, and then ‘ZOTU’ (zero-radius Operational Taxonomic Unit) representing unique bacteria was generated. The taxonomy of the denoised sequences was mapped against the RDP (Ribosomal Database Project) database (http://rdp.cme.msu.edu/) using the SINTAX algorithm. The ZOTU table was then rarefied to the minimum sequence depth for α- and β-diversity analyses.

### Analysis of generalists and specialists in root compartments

The habitat generalists and specialists of bacterial ZOTUs were identified based on specialization values with permutation algorithms (1000 permutations) by using the “Eco-IUtis” (https://github.com/GuillemSalazar/EcolUtils) [49]. When the habitat specialization values exceed the upper 95% confidence interval or fall below the lower 95% confidence interval of 1000 permutations, they were identified as habitat generalists or specialists, respectively. The Levin’s niche breadth index was calculated separately for generalists and specialists with the ‘spaa’ R package.

### Co-occurrence network construction and calculation of topological features

To reduce rare ASVs in the dataset, only ASVs that contained more than 20 reads and were present in more than 30% of each of the compartment samples were retained for co-occurrence network construction. Spearman correlation matrices were computed using the R ‘WGCNA’ package [50], and Benjamini-Hochberg false discovery rate (FDR) was used to adjust the *P*-values in the correction. Only robust correlations with Spearman’s correlation coefficients (ρ) > 0.75 or < -0.75 with corresponding *P*-value < 0.01 were included into the network analyses [51].

We then calculated the node-level topological features, including degree (the number of correlations a node has), betweenness centrality (the number of shortest paths going through a node) and eigenvector centrality (the centrality of the nodes to which it is connected) using the igraph package. High values of these topological features indicate that a node occupies a central position in the network, while low values suggest that it occupies a peripheral position [50, 51].

### Null model and neutral model analysis

The Sloan neutral model was applied to evaluate the effects of neutral processes (e.g. random dispersal and ecological drift) on the assembly of bacterial community in each root-associated compartment [52, 53]. In this model, a single free parameter *m* represents the migration rate and the parameter *R^2^* indicates the overall fit to the neutral model [52]. To assess the assembly processes of habitat generalists and specialists, null model analysis was performed using iCAMP (Additional files, [54]).

### Statistical analyses

All statistical analyses were conducted using R version 3.6.0. Differences in α-diversity, plant properties, relative expression of root transporter genes, Cd, Zn and nutrient concentrations among more than two groups were determined using ANOVA with Tukey’s HSD *post hoc* test, and difference in microbial effect between two groups was conducted with independent *t* test. Significant differences in the relative abundance of the bacterial taxa and topological parameters between two groups and more than two groups were analyzed by Wilcoxon test and Kruskal-Wallis test with *post hoc* by Dunn test. PCoA analyses were conducted using the pcoa() function, PERMANOVA was conducted using the adonis() function. Differential abundance analyses were performed with the DESeq2 package [55]. Phylogenetic tree of habitat generalists and core microbes were generated using ‘FastTree’ (v.2.1.10) with GTR (general, time-reversible) nucleotide substitution model, and were visualized and edited in the Interactive Tree Of Life [56]. Random Forest (RF) regression analysis was conducted using the ‘randomForest’ package [57]. The importance of each predictor variable is determined by evaluating the decrease in prediction accuracy (that is, increase in the MSE (mean squared error) of variables: high MSE% values indicate more important variables) when the data for that predictor is randomly permuted. Phylogenetic Investigation of Communities by Reconstruction of Unobserved States 2 (PICRUSt2) was conducted to predict functional profiles for representative 16S rRNA sequences from ASVs according to the Kyoto Encyclopedia of Genes and Genomes (KEGG) database [58].

## Results

### Host traits of distinct Cd-accumulator plants

*S. nigrum* grown in HP and SP soils had a largest shoot and root biomass, followed by *S. alfredii* and *B. juncea*, which had a comparable plant biomass that was significantly higher than that of non-hyperaccumulator *S. alfredii* (*P* < 0.05; Figure 1a,b). Shoot Cd concentrations were significantly higher in *S. alfredii* than in *S. nigrum*, followed by *B. juncea* and non-hyperaccumulator *S. alfredii* growing in these two soils had values that were markedly lower than the former two (*P* < 0.05; Figure 1c). Shoot Zn concentrations exhibited a trend similar to shoot Cd concentrations in different plants, with the exception of the significantly higher shoot Zn concentration in *B. juncea* than in non-hyperaccumulator *S. alfredii* (*P* < 0.05; Figure 1d).

**Figure 1.**
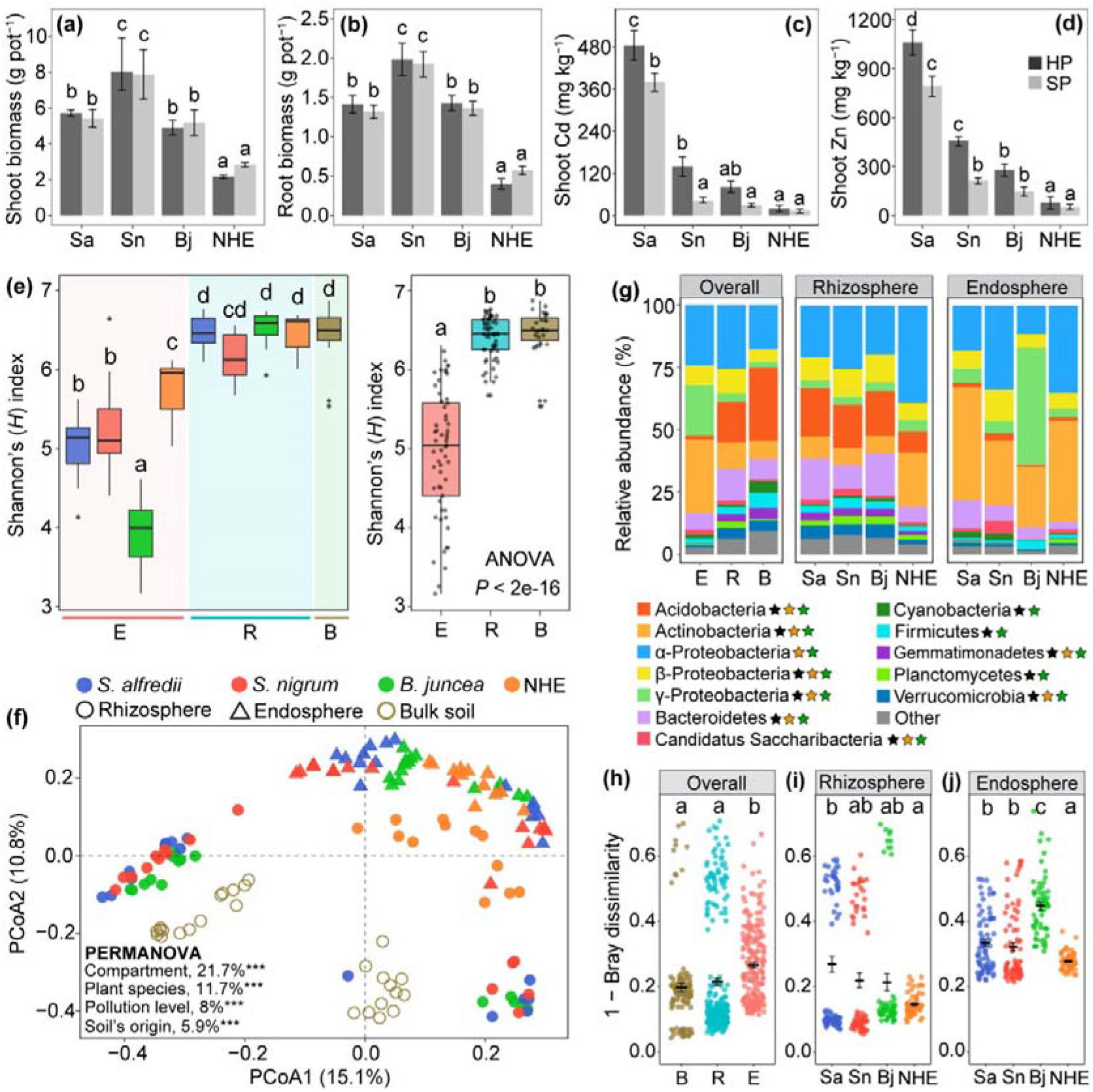
Plant traits, the diversity and composition of bacterial communities inhabiting bulk soil, rhizosphere and root endosphere associated with different plants. (**a**) Shoot biomass, (**b**) root biomass, shoot Cd concentrations (**c**) and shoot Zn concentrations (**d**) of four plants grown in HP and SP soils. (**e**) Shannon diversity index of bacterial communities inhabiting root compartments of distinct Cd-accumulators. (**f**) PCoA plot for all greenhouse root-associated and bulk soil samples generated using the Bray–Curtis distance. Significance of the biotic and abiotic driving factors on bacterial community dissimilarities was calculated based on PERMANOVA, *** *P* < 0.001. (**g**) The relative abundance of the 12 most abundant bacterial phyla/classes were significantly affected (Kruskal-Wallis; *P_adj_*< 0.05) by compartment (black star) and plant species (yellow star, rhizosphere; green star, endosphere). Pairwise distances between each type of soil within each compartment (**h**) and within each plant species for rhizosphere (**i**) and endosphere (**j**) compartment. Different letters above the bar plots, box plots and scatter diagrams indicate significant differences based on Tukey’s HSD test. Abbreviations: Bulk soil, B; Rhizosphere, R; Endosphere, E.

### The influence of plant-specific and environmental factors on the Cd-accumulator root-associated microbiota

The levels of bacterial α-diversity, estimated with the Shannon’s diversity, were highly determined by compartment type (*P* = 10^-16^, 52.5% of variance; Additional Table S2), followed by plant interspecific variation (*P* = 10^-16^, 20.3% of variance) and the interactions of these two factors (*P* = 10^-16^, 13.1% of variance; Additional Table S2). Consistent with other root microbiome studies [9, 13, 45], α-diversity decreased sequentially from the soil to the endosphere, although rhizosphere communities displayed a comparable α-diversity to bulk soil (Figure 1e). We found significant effects of root compartment (*R^2^* = 0.22, *P* < 0.001, PERMANOVA) and plant species (*R^2^* = 0.12, *P* < 0.001, PERMANOVA; Additional Table S2) on root-associated bacterial communities, when using the Bray–Curtis dissimilarity metric. Root-associated microbiomes were significantly different and clustered distinctly from those in the bulk soils (Figure 1f; Additional Table S2). Similar patterns were also observed when using other dissimilarity metrics (Additional Table S2). We noticed that the endosphere microbiota diversity and composition were more significantly affected by plant species compared to the rhizosphere counterparts, and the effect size was highly consistent across distinct soils (Additional Figure S1). Although soil’s geographical origin (*R^2^* = 0.002, *P* < 0.001; Additional Table S2) and pollution level (*R^2^* = 0.011, *P* < 0.001) had a marginal effect on the total bacterial diversity, they did affect the community composition (*R^2^* = 0.059, *P* < 0.001). The effects of soil’s geographical origin and pollution degree decreased from bulk soil to rhizosphere to endosphere (Additional Table S2). Our results suggest that bacterial communities living inside the roots of Cd-accumulators were less affected by soil changes than root external consortia.

We found that the root microbiome composition was significantly affected by the plant growth environment. Bacterial communities established under different growing conditions differed significantly in α-diversity (*P* = 0.021, ANOVA) and composition (*R^2^* = 0.15, *P* < 0.001, PERMANOVA; Additional Table S3 and Figure S2). PERMANOVA analysis showed that soil pollution level (*R^2^* = 0.17, *P* < 0.001) and plant growing environment (*R^2^* = 0.15, *P* < 0.001) explained more variability in bacterial community composition than did compartment (*R^2^* = 0.11, *P* < 0.001). The growing environment effect decreased from bulk soil (*R^2^*= 0.41, *P* < 0.001, PERMANOVA) over rhizosphere soil (*R^2^* = 0.39, *P* < 0.001, PERMANOVA) to endosphere (*R^2^* = 0.24, *P* < 0.001, PERMANOVA), and soil contamination showed an analogous pattern of influence (Additional Table S3).

### Convergence in root microbiota composition of Cd-accumulator plants

Across the three soils, the distribution patterns of the relative abundance of bacterial communities at phylum and order ranks was highly similar across the transition from bulk soil to endosphere (Additional Figure S4), suggesting a predictable and conserved assembly of bacterial root microbiota. Soil type-dependent differences in bacterial communities at phylum rank converged significantly towards more soil type-independent community profiles from bulk soil to root endosphere (Additional Figure S3a and Table S4). The percentage of shared abundant bacterial ZOTUs (>20 sequences) is significantly higher across endosphere and rhizosphere than across bulk soil samples (*P* < 0.05; Additional Figure S5). Similarly, the Bray-Curtis dissimilarities between tested soils largely decreased from root exterior to root endosphere (Figure 1h-i). These results strongly suggest that the root environment of Cd-accumulators drives convergence in bacterial community composition across distinct soils.

### Distinct ecological assembly processes driving generalist and specialist microbiota

A total of 812, 1507 and 758 bacterial ASVs were identified as generalist in bulk soil, rhizosphere and endosphere, respectively (Figure 2a,b). The distinction between specialists and generalists was confirmed by using niche breadth estimates, as expected, the generalists had a wider niche breadth than specialists (Figure 2b). The generalist microbiota within the root compartments accounted for 0.95-3.5% of the total community abundance in the different compartments. The assemblage of generalists was closely related to the total microbiota, as evidenced by high similarity of β-diversity between the identified generalist populations and the total microbiota across different compartments (Mantel’s *r* of 0.79 to 0.92, *P* < 0.001). Moreover, the taxonomic composition and predictive metagenome functions of the generalists also differed significantly between compartments (Figure 2c and Additional Figure S6).

**Figure 2.**
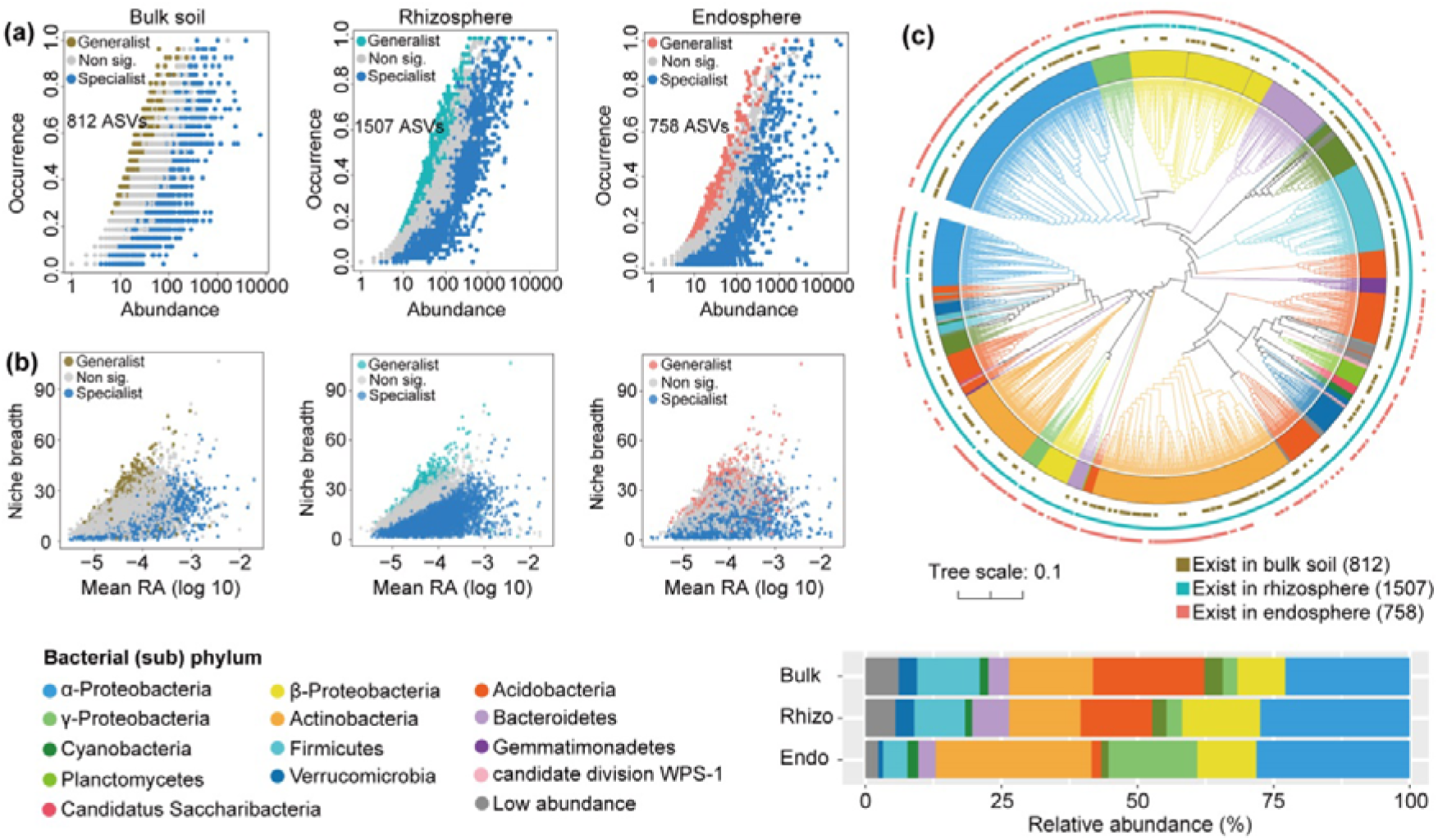
Habitat specialization, phylogenetic classification and potential ecological functions of generalist microbiota inhabiting different root-associated compartments. (**a**) Abundance-occupancy distributions were used to identify the generalists and specialists within each compartment. (**b**) Niche breadth of the generalists and specialists in the bulk soil, the rhizosphere and the endosphere, the relative abundance is shown as a log10 scaled value. (**c**) Maximum likelihood phylogenetic tree and taxonomic composition of the ZOTUs of generalist microbiota.

The neutral and null models were combined to examine the ecological assembly processes between generalists and specialists in each soil-root associated compartment. Neutral processes explained 41.8%, 18% and 61% of the ZOTUs distribution of bulk soil, rhizosphere and endosphere generalists, respectively, with 74.5%, 31.6% and 64.3% ASVs could be predicted as neutrally distributed ASVs (Figure 3a,b and Additional Figure S7a). Although the goodness of fit was similar for rhizosphere and endosphere specialists, more endosphere specialist ASVs (69.2%) were predicted as neutrally distributed compared with their rhizosphere counterparts (54.3%; Figure 3c,d). The generalists in each compartment had substantially higher migration rates (*m* values) than the specialists, confirming that generalists are less constrained by dispersal limitation (Figure 3a-d and Additional Figure S7a,b). Furthermore, we deeply elucidated the assembly processes based on the null model. A deterministic process, in particular, variable selection was the most important assembly process of endosphere, rhizosphere and bulk soil generalists, with their relative contributions of 53.7%, 54.8% and 77.2%, respectively (Figure 3e and Additional Figure S7c). Variable selection contributed more to endosphere specialists (64.8%) than to rhizosphere specialists (41.5%), and dispersal limitation exhibited an opposite trend (Figure 3e). The endosphere generalists (34%) were more influenced by dispersal limitation compared to the rhizosphere generalists (21.1%). Consequently, deterministic (54.8% vs. 64.8%) and differentiating (75.9% vs. 97.1%) processes determined the assembly of rhizosphere generalists and endophytic specialists, and stochastic (58.5% vs. 53.7%) and differentiating (84.5% vs. 87.7%) processes governed rhizosphere specialist and endophytic generalist assembly (Figure 3f).

**Figure 3.**
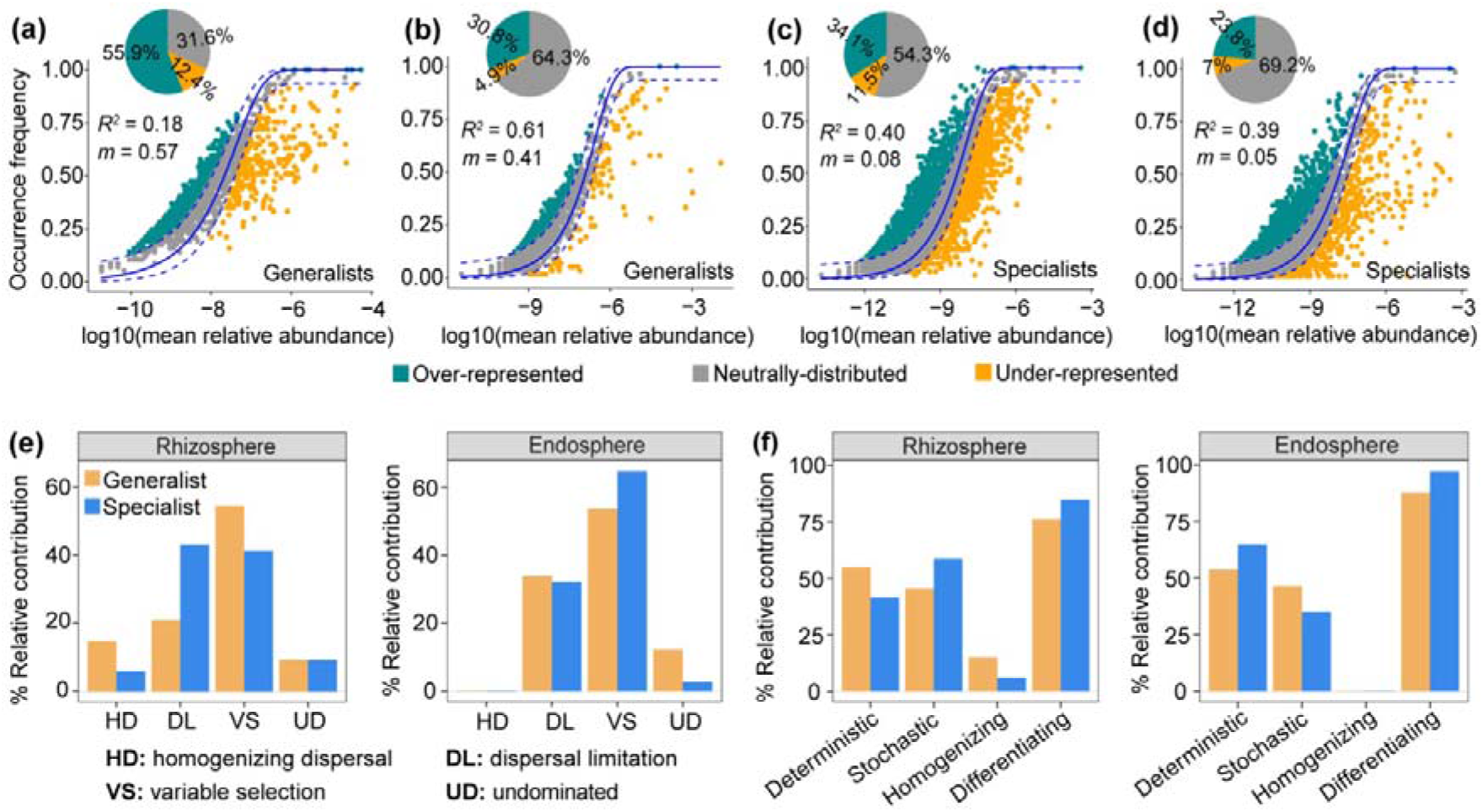
Community assembly patterns of bacterial sub-communities associated with three Cd-accumulators and a non-hyperaccumulator plant grown under greenhouse conditions. Neutral model applied to assess the effects of random dispersal and ecological drift on the assembly of rhizosphere (**a**) and endosphere generalists (**b**), and rhizosphere (**c**) and endosphere specialists (**d**). The solid blue lines indicate the best-fit to the neutral model, and dashed blue lines represent 95% confidence intervals around the model prediction. The pie charts depict the ratio of the over-represented, neutrally distributed and under-represented OTUs in habitat generalists–specialists. (**e**) Null model applied to assess the effects of variable selection, dispersal limitation, homogenizing dispersal and undominated of the habitat generalists and specialists. (**f**) The relative contribution of deterministic, stochastic, homogenizing and differentiating processes of the habitat generalists and specialists. Deterministic = Variable selection; Stochastic = Dispersal limitation + Homogenizing dispersal; Homogenizing = Homogenizing dispersal; Differentiating = Variable selection + Dispersal limitation.

### Co-occurrence networks of generalists and specialists in distinct root compartments

The rhizosphere network presented the highest number of nodes (2378), edges (60710) and connections per node (average degree: 41.56) in comparison with the bulk soil and endosphere networks (Additional Table S5). However, the network of endophytic communities showed a highest level of modular structure and average clustering coefficient but a lowest average path length relative to the networks of other communities. The empirical network of bacterial communities within each compartment exhibited higher values of average clustering coefficient, average path length and modularity than those of the respective Erdós-Rényi random network, indicating that the bacterial co-occurrence network displayed a small-world pattern and modular structure (Additional Table S5).

Among all nodes within the network of each compartment, specialists had a substantially higher number of nodes and edges than generalists. The inner associations among each sub-community were 1316/11738, 2564/33508, and 705/4356 for generalists/specialists in the bulk soil, the rhizosphere and the endosphere networks, respectively (Additional Figure S8a-c). We also characterized the node-level topological features of different sub-communities within each compartment (Additional Figure S8d-f and S9). The degree, betweenness centrality and eigenvector centrality were significantly higher (*P* < 0.005) in the specialists than in the generalists (except for eigenvector centrality in endosphere network) (Additional Figure S9). Only the eigenvector centrality of generalist networks significantly increased from bulk soil to rhizosphere and to endosphere (Additional Figure S8g-i). These three topological parameters of specialist networks were highest in the rhizosphere and lowest in the endosphere (Additional Figure S8g-i). When the network was present in a modular structure, most nodes were distributed in six modules, which accounted for 81.2%, 89.5% and 60.1% of the total nodes in bulk soil, rhizosphere and endosphere, respectively (Additional Figure S8d-f,j).

### Generalist core microbiota and Cd-accumulatorts exhibit stable associations

There are 24 and 9 ZOTUs identified as members of the core rhizosphere and endosphere microbiota (defined as generalist ZOTUs detected in at least 90% of the rhizosphere or endosphere samples), with seven ZOTUs overlapping between these two compartments (Figure 4a). The rhizosphere core microbiota represented 2% of the bacterial reads and 35.7-86.5% of the relative abundance of the generalist microbiota. These 24 ZOTUs, which belong to diverse genera (for example, *Sphingomonas*, *Bradyrhizobium*, *Devosia*, *Mesorhizobium*, *Bacillus*, *Neobacillus*, *Curvibacter* and *Burkholderia*), and 11 of them were significantly enriched in rhizosphere relative to bulk soil (*P*_adj_ < 0.05, Additional Table S6). The endosphere core taxa accounted for 0.6% of the bacterial reads and 16.2-66% of the generalist bacterial communities. At genus level, the nine endosphere cores were affiliated to *Bradyrhizobium*, *Devosia*, *Mesorhizobium*, *Bosea*, *Acidovorax*, *Bacillus* and *Sphingomonas*, with seven of them being more abundant in the endosphere than in the rhizosphere or bulk soil (*P*_adj_ < 0.05; Figure 4b and Additional Table S6). Interestingly, six of them were more abundant in *S. alfredii* and *S. nigrum* than in *B. juncea* and non-hyperaccumulating *S. alfredii* (Wilcoxon test; *P* < 0.05; Figure 4c), with the former two having a stronger ability to accumulate Cd/Zn than the latter two. Such an enrichment pattern is true for only seven core rhizosphere ZOTUs (Additional Figure S10). In addition, 67% of rhizosphere and 56% of endosphere core ZOTUs were predicted above the neutral model partition, suggesting that majority of core taxa were deterministically-selected by the root environment (Additional Figure S11). Further inspection of the generalist microbiota revealed a high level of conservation for some core members, with 54% of rhizosphere and 67% of endosphere core microbes being consistently detected in the same soils under field and controlled conditions (Figure 4d-g).

**Figure 4.**
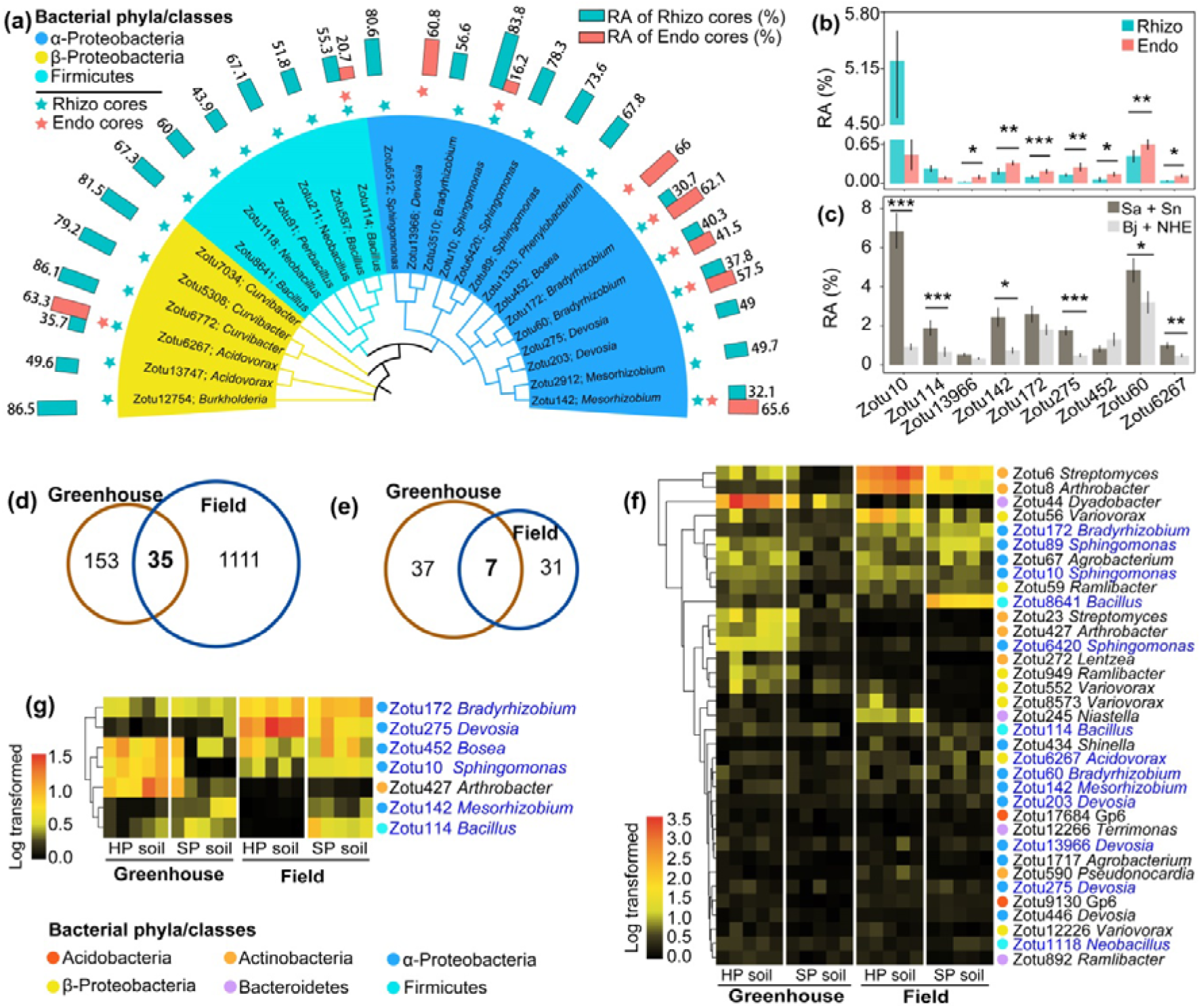
Characterization and stability of the generalist core microbiota inhabiting the rhizosphere (Rhizo) and endosphere (Endo). (**a**) Maximum likelihood phylogenetic tree for the generalist core ZOTUs. The tree clades were colored according to the taxonomic affiliation of each core ZOTUs. The bars outside the tree represent the read proportion of a core ZOTU to the rhizosphere or endosphere generalist microbiota. The sky blue and red asterisks denote the rhizosphere and endosphere generalist core taxa, respectively. (**b**) Comparison of the relative abundance (RA) of the generalist core ZOTUs between compartments using DESeq2, only the RA was significantly higher in endosphere than in rhizosphere are labeled, * *P* < 0.05, ** *P* < 0.01, *** *P* < 0.001. (**c**) Comparison of the RA of the generalist core endosphere ZOTUs of the two plants exhibiting stronger Cd/Zn accumulation ability (*S. alfredii* and *S. nigrum*) with the two plants exhibiting weaker ability (*B. juncea* and non-hyperaccumulator *S. alfredii*) using Wilcoxon test. * *P* < 0.05, ** *P* < 0.01, *** *P* < 0.001. Venn diagrams of the rhizosphere (**d**) and endosphere (**e**) bacterial populations overlapped between the greenhouse and field conditions. Heatmap showing the RA (log-scaled) and taxonomic classification of the rhizosphere (**f**) and endophytic (**g**) ZOTUs shared under greenhouse and field conditions. The generalist core taxa are highlighted in dark blue.

### Role of generalists core microbes in maintaining network stability and predicting plant functional traits

The network of generalist microbiota showed that the degree and betweenness centrality in the endosphere network were significantly higher for generalist core taxa than for non-core taxa, while the degree of the bulk soil network exhibited an opposite trend (Additional Figure S12a-c). Spearman correlation analysis showed that the relative abundance of most generalist core microbes (22/24) inhabiting the rhizosphere was positively correlated with one or numerous plant characteristics (*P* < 0.05), such as shoot and root biomass, and plant accumulation of Cd, Zn, Cu, N and P (Additional Figure S12d). For example, the relative abundance of 11 ZOTUs belonging to *Sphingomonas*, *Bacillus*, *Neobacillus*, *Mesorhizobium*, *Peribacillus* and *Devosia* was highly correlated with shoot and root biomass (Spearman’s ρ = 0.30–0.57, *P* < 0.05; Additional Figure S12d). The relative abundance of 11 different ZOTUs was positively correlated to plant shoot N and P accumulation (*P* < 0.05). Additionally, we noticed that the prevalence of six core rhizosphere ZOTUs exhibited a significant positive correlation with shoot Cd accumulation, while eight ZOTUs showed a significant (*P* < 0.05) positive correlation with shoot Zn accumulation. With respect to the endosphere cores, the relative abundance of a ZOTU belonging to genus *Sphingomonas* was highly positively correlated to shoot and root biomass (*P* < 0.001; Additional Figure S12e). The abundance of three other ZOTUs was positively related to shoot Cd accumulation, and the abundance of another three ZOTUs was significantly correlated to shoot Zn accumulation (*P* < 0.05). Furthermore, the random forest model results suggested that the generalist core populations explained more variation in plant biomass (61% vs. 36%) and shoot N and P accumulation (27% vs. 12%), but less variation in plant metal accumulation (10% vs. 19%) when inhabiting the rhizosphere than the endosphere (Additional Figure S12d,e).

### Isolation of core bacteria and their capability for plant growth promotion and Cd and nutrient accumulation

Pot experiments were conducted to assess the capability of the core rhizosphere and endosphere microbes to promote *S. alfredii* growth, Cd and nutrient accumulation. We found that SynCom comprising 14 core endophytes (Figure 5a) significantly increased the relative expression of transporter genes of *SaIRT*1, *SaHMA*2, *SaHMA*3, *SaNRAMP*3, *SaNRAMP*6 and *SaZIP*1 as compared with Cd and heat killed SynCom treatments (*P* < 0.05; Figure 5b). Inoculation of the endophytic SynCom significantly enhanced plant shoot biomass, shoot and root Cd accumulation, total root length and root surface area (*P* < 0.05; Figure 5b-f,g-l). Endophytic SynCom inoculation resulted in a markedly higher microbial effect than inoculation of heat killed SynCom (*P* < 0.05; Figure 5m). In addition, we observed that inoculation of the SynCom comprising 17 core rhizosphere bacterial isolates led to significance increase of shoot biomass, total root length, shoot and root Cd accumulation but not the concentration of Cd per unit of plant tissues compared with other treatments (*p* < 0.05; Additional Figure S13b-j). Core rhizosphere SynCom significantly increased plant shoot nitrogen and phosphorus contents and microbial effect of Cd, total N and total P (*P* < 0.05; Additional Figure S13k,l and S14). Finally, soil available nutrients of NH_4_^+^, NO ^-^ and available phosphorus were significantly higher in SynCom-inoculated soil than those in soils subjected to other treatments (*P* < 0.05; Additional Figure S13m-o).

**Figure 5.**
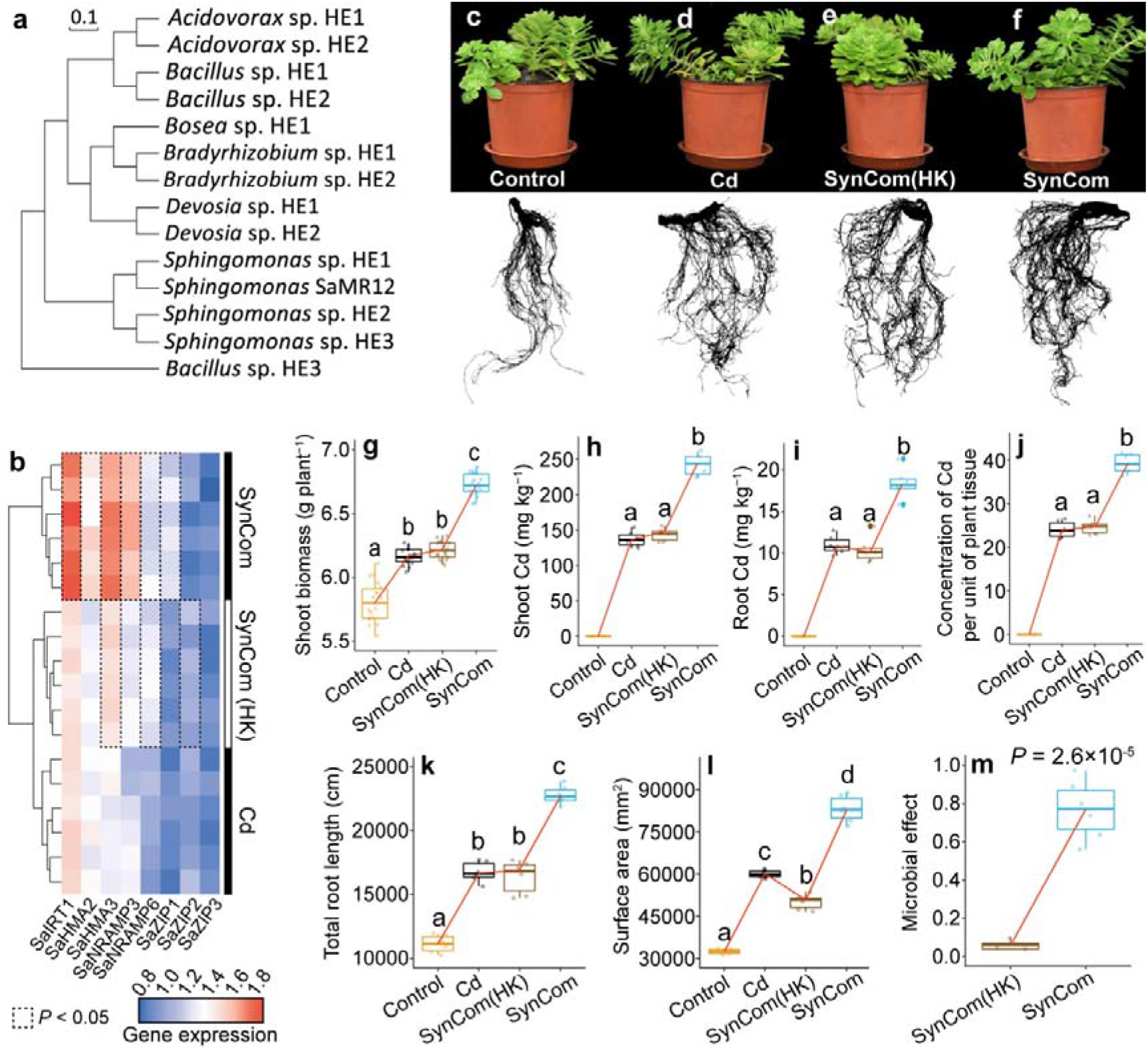
SynCom comprising the isolates of core endophytic bacteria promotes *S. alfredii* growth, facilitates Cd accumulation. Phylogenetic tree of 14 endophytic bacterial isolates used to establish the SynCom, based on the distance values calculated from average nucleotide identity of their full-length 16S rRNA gene with the maximum likelihood method (**a**). Relative expression of the selected metal transporter genes within *S. alfredii* roots (**b**). Comparisons with *P*-value < 0.05 are contoured in black dotted line. Growth status of *S. alfredii* cultivated in different treatments (**c-f**). Effects of Cd, SynCom, heat-killed SynCom [SynCom(HK)] and control (no Cd and SynCom added) treatments on shoot dry weight (**g**), concentrations of Cd in shoot (**h**) and root (**i**) and per unit of plant tissue (**j**), total root length (**k**), root surface area (**l**) and microbial effect on plant Cd accumulation (**m**). Values are mean ± s.d. (shown as error bars). Different letters above the box plots indicate significant differences according to Tukey’s HSD test at *P* < 0.05.

## Discussion

Our work provides valuable evidence on host selection and the increasing convergence in root microbiota composition along the soil-root continuum in Cd-accumulators. We identified a group of generalist core microbes as the host-specific and conserved signatures in the root-associated microbiota between the Cd-accumulators, and the relative abundance of these core microbes increased with the increasing ability of plant to accumulate Cd and Zn. These result hinted that the establishment of core microbiota was more driven by shared environmental and physiological traits of the host plants than by environmental parameters. Co-occurrence network and random-forest analyses identified that the core microbiota inhabiting different root compartments played differential roles in community stability and plant functional traits. Inoculation of hyperaccumulator *S. alfredii* with SynCom comprising core microbes demonstrated that rhizosphere cores played important roles in plant growth promotion and nutrient acquisition, while endosphere core were more important in root development and plant Cd accumulation. Therefore, the host plants with similar functional traits tend to have the potential to recruit the conserved microbiota that reciprocally provides benefits to plant characteristics. This study brings us closer to developing core microbiome-based strategies to improve plant productivity and optimize phytoremediation systems through host genetics.

### Major and core microbial players highlight convergence across the Cd-accumulators

Our results demonstrate that the roots of distinct Cd-accumulators provide a niche for the assembly of a conserved bacterial consortium at a broad environmental scale, despite extensive differences in the indigenous soil microbiota and physicochemical properties between soils [59]. While the rhizosphere and endosphere bacterial communities vary largely between plant species, a set of bacterial assemblages have been identified as the conserved microbiota across different Cd-accumulators, and probably explains the similarity in bacterial community composition observed across microbiome studies [60, 61]. For example, *Bradyrhizobium*, *Burkholderia* and *Acidovorax* (Zotu3510, Zotu172, Zotu60, Zotu12754, Zotu13747 and Zotu6267 in this study) were the three most dominant of the 47 widespread genera in the roots of 31 diverse plant species across a 10-km transect in Australia [60] and of the 13 ubiquitous OTUs in the roots of *Arabidopsis thaliana* across 17 European sites [62]. In addition, our observation that average 62% of the core microbes from the root compartments were above the neutral model prediction could illustrate that a majority of core taxa are positively selected by the plant environment [26]. Among these cores, 46% and 78% of them were significantly enriched in the rhizosphere and endosphere, respectively, as compared with the bulk soil. This suggests not only that these core members were able to persistently colonize the plant roots but also that they reached a higher abundance within the root environment, implying some degree of adaptation to host niches [17]. Besides, a large part of the core microbes (∼61% ZOTUs) from the greenhouse condition overlapped with those detected in the field, further suggesting a plant-driven selection for core microbiome. Overall, these information, along with the remarkable phylogenetic diversity among these core taxa, indicative of convergent evolution and metabolic adaptation to the root niche of multiple Cd-accumulators in phylogenetically distant bacterial lineages [62]. Therefore, our study provides evidence suggesting that the establishment of core microbiota in the roots of multiple metal accumulators was primarily driven by shared environmental and physiological traits of the host plants, rather than by environmental parameters [63]. These findings could partially support the view that host-associated bacterial community assembly based on functional profiles rather than species, such as the features related to a host-associated lifestyle [64].

### Differential assembly processes dominated the habitat generalists and specialists

In this study, neutral processes explained more variation in assembly of habitat generalist and specialist subcommunities in the endosphere compared to the rhizosphere. This indicates the stochastic nature of microbial endosphere colonization, as reported in previous studies [65]. Through null model and neutral model analyses, we found that habitat generalists and specialists were differentially driven by deterministic and stochastic processes, consistent with ecological theory demonstrating that habitat disturbances (e.g. heavy metals and plant physiological perturbations in this study) differentially modulate the distributions of habitat generalist and specialist species [66]. Here, deterministic processes were more important for the assembly of specialists in endosphere than in rhizosphere. Habitat specialists with narrower niches and lower environmental tolerance are considered to be particularly sensitive to environmental disturbance and variability possibly because they are metabolically inactive [25], and thus are more likely to be shaped by deterministic processes [67]. Since endophytes are expected to have a more intimate relationship with their hosts than epiphytes [68], they may have evolved to specialize in a particular habitat within the root and exhibit specific environmental fitness [69]. This makes stochastic processes (i.e. dispersal limitation) have a greater impact on the assembly of endosphere generalists than rhizosphere generalists. Habitat generalists are metabolically more flexible and feature higher resistance against changing conditions than specialists, and thus habitat disturbance favors generalists [25, 66]. Previous studies have shown that higher concentrations of heavy metals in metal-accumulators than in the soil (4-34 times higher in SP soil) could drive endophytes to more adapt to such environmental conditions and plant physiological changes than epiphytes [70, 71]. This microbial characteristic may thus allow endophytes to be less affected by deterministic processes.

### Co-occurrence networks of generalists and specialists and the associations of the generalist core microbes with plant traits

In ecology, modular structure may suggest niche overlap, phylogenetic clustering of closely related species and even a key unit of species co-evolution [72, 73]. A higher network clustering coefficient implies a greater number of edges within clusters than between clusters [74], implying higher associations between bacteria in similar niches within roots. Average path length, defined as the average number of steps along the shortest paths between each node, is a measure of efficiency of a network [75]. Therefore, modularity combined with a short average path length may represent a faster response and higher resistance of the community to environmental changes [74]. Endophytic microbiota, which are subject to stronger plant selective pressures than the rhizosphere and bulk soil communities, are expected to form a less complex network [76]. However, the endosphere co-occurrence networks exhibited the highest level of modularity and clustering coefficient, but the lowest average path length. These suggested that the endosphere microbiota was more resistant than the rhizosphere and bulk soil microbiota to such changes. Furthermore, the larger network sizes and higher complexity were observed for specialists than for generalists, indicating that the former may interact more frequently with each other than do the latter [77]. Similarly, the significantly higher values of betweenness centrality and eigenvector centrality for specialists than for generalists may suggest that specialists are more often located in central positions within the network than generalists (Additional Figure S8 and S9), and thus had a higher likelihood of becoming ‘hub microorganisms’ [78]. Hub species play a crucial role in organizing the plant microbiota as a network through biotic interactions and selective assembly, and thereby pay a key role in orchestrating host–microbiome interactions and maintaining ecosystem stability [7].

The core plant microbiota generally refer to microbial communities that are systematically associated with a given host and established through a long-lasting evolutionary mechanism of selective pressures that shape microbiome assembly [15]. In this study, we identified a group of core rhizosphere and endophytic bacteria consisting of generalist species that were conserved across different Cd-accumulators grown in controlled and field environments. Among the rhizosphere cores, members of *Sphingomonas* and *Burkholderia* were also the core microbes that were identified by a different method in our recent study [40]. This indicates the selective enrichment of these cores driven by plant roots and that the structure of the core microbiota is independent of the different analytical methods used. Multiple members affiliated with these core microbes, such as *Sphingomonas*, *Burkholderia*, *Bradyrhizobium* and *Mesorhizobium*, were found to be positively related to different plant functional traits, including plant biomass, nutrient and Cd/Zn accumulation, providing supports for the notion that core microbes probably contribute to plant growth and performance of plant functional traits [19, 28]. A notable point is the close links of core microbes affiliated with *Bradyrhizobium* and *Mesorhizobium* to plant Cd and Zn accumulation. These two bacterial groups are well-known root-nodulating bacteria and provide their hosts with biologically fixed nitrogen [79], however, the role of nitrogen-fixing microorganisms in phytoremediation systems has rarely been recognized [80].

The existence of common core microbes with beneficial functions in different Cd-accumulators indicates that a highly conserved and coevolutionary core root microbiota may exist and play an essential role in maintaining holobiont fitness [1, 27], which should simplify the identification of microbes as targets for future investigation into their role in plant growth and plant characteristics. The generalist core members inhabiting the endosphere tend to occupy central positions with remarkably higher connectivity and centrality within the networks as compared with other generalist members (non-core). In network analysis, centrality indicates the most important nodes, which can be interpreted as key taxa within a connected community [81]. Therefore, the close interaction of generalist core microbes inhabiting the endosphere may have a greater potential to facilitate microbiome diversity and stability, and to contribute to host fitness and functional traits. Moreover, generalist core microbes inhabiting the rhizosphere were more closely linked to plant phenotypic traits, such as plant biomass and aboveground N and P accumulation, and act as more important predictors of these traits than those inhabiting root endosphere. Previous studies revealed that plant physiological and morphological traits are usually linked to the strategies of plant resource acquisition, thus highlighting the ecological importance of microbial processes and interactions in the rhizosphere for plant resource consumption and turnover [82]. This is supported, at least in part, by our result that α- and β-Proteobacteria being dominant members (75%) of the core rhizosphere microbiota, as members of these two Proteobacterial groups are generally fast-growing *r*-strategists with the ability to utilize a variety of carbon resources [83], which should facilitate nutrient turnover. In comparison, the presence of core microbes in the root endosphere is likely the result of a long and intimate history of coevolution between host plant and its microbiome, which becomes highly adapted to and dependent on host life without necessarily being directly beneficial. Other evidences supporting this hypothesis were that 67% core microbes were significantly more abundant in the two plants with stronger Cd/Zn accumulation ability (*S. alfredii* and *S. nigrum*) than other two plants with weaker Cd/Zn accumulation ability (*B. juncea* and non-hyperaccumulating *S. alfredii*). These may indicate that the higher abundance and occurrence of core microbiome may aid plant Cd/Zn accumulation.

### Variability and stability of Cd-accumulator root-associated bacterial communities

Our results found that the observed species effect size remained highly stable across the three soils, suggesting that plant-related factors driving root microbiome assembly were predictable and less confounded by variation in soil and indigenous microbiota. Within each compartment, the diversity and composition of the endosphere microbiota were defined more strongly by plant species than by environmental factors, while this case was not true for the rhizosphere microbiota. This supports the idea that host-selective forces on the microbiome grow in magnitude with increased plant-microbe intimacy, as was reported in studies of 18 grass species and 30 angiosperm plants [84, 85]. Together, these results highlight the existence of coevolution between host plant and its microbiota, leading to filtered subsets of soil microbiota with niche-specific adaptation ability [86].

## Conclusions

Our study demonstrates that Cd-accumulator microbiome assembly along the soil-root continuum is predominantly driven by host selection, rather than by environmental parameters.

We observed that the existence of a largely conserved and taxonomically narrow root core microbiota between the tested Cd-accumulators in controlled and field environments that are functionally important for plant characteristics suggests common selective pressures governing core microbiota assembly, possibly via shared environmental and physiological traits of plants. Therefore, it is possible that the conserved core microbiota between different plants represents a standing reservoir of retrievable host plant functional traits that can be exploited to improve plant growth and phenotypic characteristics.

## Supplementary Information

**Additional file 1: Figure S1.** PCoA plot highlighting a stable influence of plant interspecific variation on the root-associated bacterial communities. **Figure S2.** Structure, diversity and relative abundance of root-associated bacterial communities established under greenhouse and field conditions. **Figure S3.** Plant growth in different soils affects root-associated bacterial community compositions. **Figure S4.** Distributions of bacterial phyla and orders across root compartments. **Figure S5.** Percentage of shared abundant ZOTUs among replicate samples in each soil across root compartments. **Figure S6.** Mean relative abundance of the 30 most abundant predicted KEGG functional orthologs of generalist microbiota. **Figure S7.** Community assembly patterns of bacterial sub-communities in the bulk soil under greenhouse conditions. **Figure S8.** Co-occurrence patterns of the habitat generalists and specialists across different root compartments. **Figure S9.** Node-level topological parameters between the habitat generalists and specialists inhabiting distinct root compartments. **Figure S10.** Comparison of the relative abundance of the 24 generalist core rhizosphere ZOTUs between host plants. **Figure S11.** Neutral model applied to assess the distribution of the generalist core ZOTUs in the rhizosphere and endosphere. **Figure S12.** Bacterial co-occurrence networks of generalist microbiota inhabiting different root compartments and linkages between generalist core microbes and plant functional traits. **Figure S13.** SynCom comprising the isolates of core rhizospheric bacteria promotes *S. alfredii* growth and improves plant nutrient status. **Figure S14.** Microbial effect of rhizospheric SynCom on plant accumulation of Cd, total nitrogen and total phosphorus.

**Additional file 2:** The materials and methods of “Characterization of experimental soils”, “Sampling of the bulk soil, rhizosphere and root endosphere compartments”, “Plant Cd accumulation, soil Cd content, and plant nitrogen and phosphorus status”, “Quantitative Real-Time PCR (qPCR)”, “Null model and neutral model analysis”. The results of “Taxonomic composition of the root microbiota composition of Cd-accumulator plants”.

**Additional file 3: Table S1.** Physiochemical parameters of the three tested soils.

**Additional file 4: Table S2.** Comparison of bacterial community diversity between different samples.

**Additional file 5: Table S3.** Comparison of bacterial community diversity across compartments, pollution degree and growing conditions.

**Additional file 6: Table S4.** The influence of compartment type and soil variation on the relative abundance of the ten most abundant bacterial phyla.

**Additional file 7: Table S5.** Topological property of co-occurrence networks within different root-associated compartments.

**Additional file 8: Table S6.** Differential abundance analysis of the generalist core ZOTUs between root compartments and bulk soil.

**Additional file 9: Table S7.** Forward and reverse primers used in quantitative PCR analysis.

## Acknowledgements

Not applicable

## Authors’ contributions

L.J. and L.T. designed and supervised the project. L.J., W.Y., Z.Y., and G.W. performed the experiment. L.J. and G.S. conducted the bioinformatics analysis. L.J., G.S. and L.T. prepared figures and tables. L.Y. and Z.H-P coordinated field experiments. G.S., L.Y. and Z.H-P suggested protocols, data analyses, and interpretation of results. L.J., L.T., and G.S. wrote and revised the manuscript. All authors have read and approved the submitted version of the manuscript.

## Funding

This research is financially supported by the National Natural Science Foundation of China (42107009, 41977107, and 42177008), the fellowship of China Postdoctoral Science Foundation (2022M712770), and the Key Research and Development Projects of Zhejiang Province, China (2022C02018, 2022C02022).

## Availability of data and materials

The raw sequencing data have been deposited in the NCBI Short Read Archive under BioProject number PRJNA691442.

## Declarations

### Ethics approval and consent to participate

Not applicable.

### Consent for publication

Not applicable.

### Competing interests

The authors declare that they have no competing interests.

### Author details

^1^Ministry of Education Key Laboratory of Environmental Remediation and Ecological Health, College of Environmental and Resource Sciences, Zhejiang University, 310058, Hangzhou China. ^2^Center for Quantitative Biology and Peking-Tsinghua Center for Life Sciences, Peking University, 100091, Beijing, China. ^3^Zhejiang Provincial Key Laboratory of Agricultural Resources and Environment, 310058, Hangzhou, China

## Notes

### Competing Interest Statement

The authors have declared no competing interest.

