## Supplementary_Information for "Host signature as major driver of root and rhizosphere core microbiomes that differently affect plant functional traits"

Number of figures: 14

Number of tables: 7

Number of pages: 28

**Materials and Methods**

**Characterization of experimental soils**

The HP and SP soils were collected from two agricultural field sites located in the southwestern part of Hangzhou, China (30°04′38.3′′N; 119°24′39.7′′E), which has been contaminated by several metal smelters that were active for more than thirty years. In contrast, the non-polluted (NP) soil was collected from a paddy field in Jian City, Jiangxi Province, China (27°21′24″N; 114°22′46″E), which is located >700 km away from Hangzhou. Soils were sampled from the uniform mineral layer at a soil depth of 0-20 cm, transported to the laboratory in plastic bags, air-dried and sieved (< 2mm) for analysis of soil properties and plant growth trials. The HP and SP soils are Eutric Fluvisol and the NP soil is Haplic Alisol. The HP soil had a pH of 6.43 and contained 12.5 mg kg^-1^ total Cd, 1.9 mg kg^-1^ DTPA-extractable Cd, 824.5 mg kg^-1^ total Zn and 22.6 mg kg^-1^ DTPA-extractable Zn, and the SP soil had a pH of 6.57 and contained 0.98 mg kg^-1^ total Cd, 0.26 mg kg^-1^ DTPA-extractable Cd, 242.7 mg kg^-1^ total Zn and 5.75 mg kg^-1^ DTPA -extractable Zn. Soil physicochemical parameters were determined using the standard methods [1]. The physicochemical parameters of the three tested soils were reported in our previous study [2] and can be found in Table S1.

**Sampling of the bulk soil, rhizosphere and root endosphere compartments**

Briefly, root systems were excavated and moderately shaken to remove loosely attached soil particles. Rhizosphere soil, which is defined as soil firmly attached to the roots (1–3 mm around the root), was detached by vigorous shaking in sterile phosphate buffer solution (PBS). The root endosphere samples were enriched for endophytic microorganisms by detaching surface-adhering microbes by sonication, following our previously described methods [3]. Bulk soil was randomly sampled from the control plots after depleting the top 1-2 cm of soil. All the plant shoot, root and soil samples were immediately frozen in liquid nitrogen and stored at -80℃ until analysis.

**Plant Cd accumulation, soil Cd content, and plant nitrogen and phosphorus status**

At harvest, the fresh root samples were immersed in 20 mM Na_2_-EDTA solution to remove the heavy metals adhering to the root surface. The root and shoot samples were rinsed with deionised water and dried at 60℃ to a constant weight. Dry root and shoot samples (~0.1 g) were grounded to powder and digested with HNO_3_-H_2_O_2_ (5:1, v/v) using microwave dissolver. Available Cd and Zn in the soil were extracted with 0.05 M DTPA solution. The Cd concentrations in the digestion and extraction solutions were determined using inductively coupled plasma mass spectrometer (ICP-MS; Agilent Technologies, CA, USA). Plant and soil total nitrogen concentrations were determined using the Kjeldahl method and plant total phosphorus concentration was determined with the molybdate-vanadate-phosphate method [4].

The net heavy metal extracted per plant (microbial effect) was calculated from the difference between the microbially treated and control samples as a percentage of the control, as previously described [5] using the following equation:

Microbial effect = (treatment value－control value)/control value

**Quantitative Real-Time PCR (qPCR)**

The relative expression of root transporter genes (Additional Table S7) was determined by qPCR. For relative expression analysis, RNA was converted into cDNA using the PrimeScript^TM^ RT reagent kit (Takara, China). The cDNA was then used in subsequent qPCR assays. *Actin* was used as internal reference gene. All of the primers are listed in Additional Table S7. The qPCR cycling program was set as previously described [6]. The relative expression of transporter genes was calculated using the 2^–△△^CT method.

**Null model and neutral model analysis**

The Sloan neutral model was applied to evaluate the effects of neutral processes (e.g. random dispersal and ecological drift) on the assembly of bacterial community in each root-associated compartment [7, 8]. This model predicts the relationships between the occurrence frequency and their relative abundance (Burns et al., 2016), where a single free parameter m is used to describe the migration rate and the parameter *R^2^* represents the overall fit to the neutral model [8]. Calculation of 95% confidence intervals around all fitting statistics was done by bootstrapping with 1000 bootstrap replicates. A higher value of *m* implies that the bacterial community is less environmentally constrained. In addition, the generalists and specialists from each compartment were subsequently separated into three partitions depending on whether they occurred more frequently than (over-represented ASVs), less frequently than (under-represented ASVs) or within (neutrally distributed ASVs) the 95% confidence interval of the neutral model predictions. Neutral model was calculated using the package ‘reltools’ in the R environment.

To assess the assembly processes of habitat generalists and specialists, null model analysis was calculated using iCAMP [9]. This method mainly classifies underlying divers of assembly processes into deterministic processes, e.g. homogeneous selection and variable (heterogeneous) selection, and stochastic processes, e.g. dispersal limitation, homogeneous dispersal and “Undominant” [10]. Briefly, this approach is based on Bray-Curtis-based Raup-Crick (RC_bray_) and beta-nearest taxon index (βNTI) to calculate the differences in taxonomic and phylogenetic diversity. According to Stegen et al. (2013) [11], the βNTI value > +2 or < -2 means significantly greater or less phylogenetic turnover than expected, respectively, indicating the predominance of deterministic processes (variable selection or homogeneous selection, respectively). In contrast, a |βNTI| ≤ 2 means non-deviance from the null distribution, indicating stochastic processes predominate. When |βNTI| <2 and RC_bray_ <- 0.95 or the |βNTI| < 2 and RC_bray_ ≥ 0.95 were identified as homogenizing dispersal and dispersal limitation respectively. The |βNTI| < 2 and RC_bray_ < 0.95 indicate the influence of the ‘Undominated’ fraction which constrains ecological drift, diversification, weak selection and/or weak dispersal.

**Results**

**Taxonomic composition of the root microbiota composition of Cd-accumulator plants**

In all samples, the bacterial community was dominated by Proteobacteria (47.3%) and Actinobacteria (20%), and to a lesser extent by Bacteroidetes (9.5%) and Acidobacteria (8.9%) (Figure 1g). The rhizosphere and endosphere were enriched for bacteria belonging to β-Proteobacteria, γ-Proteobacteria, Actinobacteria and Bacteroidetes (Kruskal-Wallis; *P* < 0.001), and depleted for Acidobacteria, Firmicutes and Gemmatimonadetes as compared to the bulk soil (Kruskal-Wallis; *P* < 0.001; Figure 1g).

**Figures**

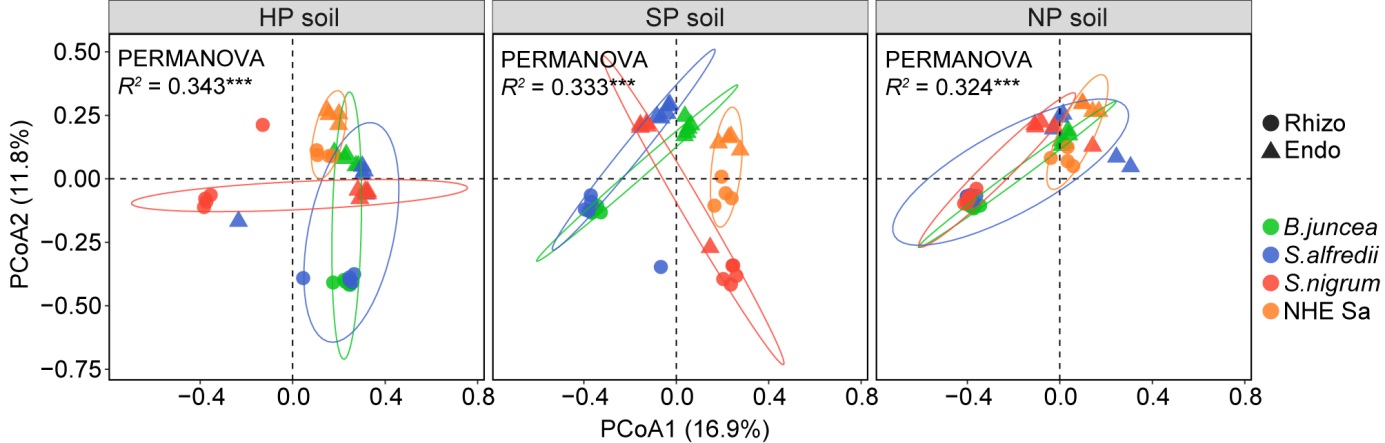
**Figure S1** PCoA plot highlighting a stable influence of plant interspecific variation on the structure of rhizosphere and root endosphere bacterial communities across different soils (HP soil, SP soil and NP soil). The *R* squared and *p*-value represent the strength and significance of plant species in community variation, respectively, as evaluated by PERMANOVA. Ellipses show the parametric smallest area around the mean that contains 95% of the probability mass for each plant species.

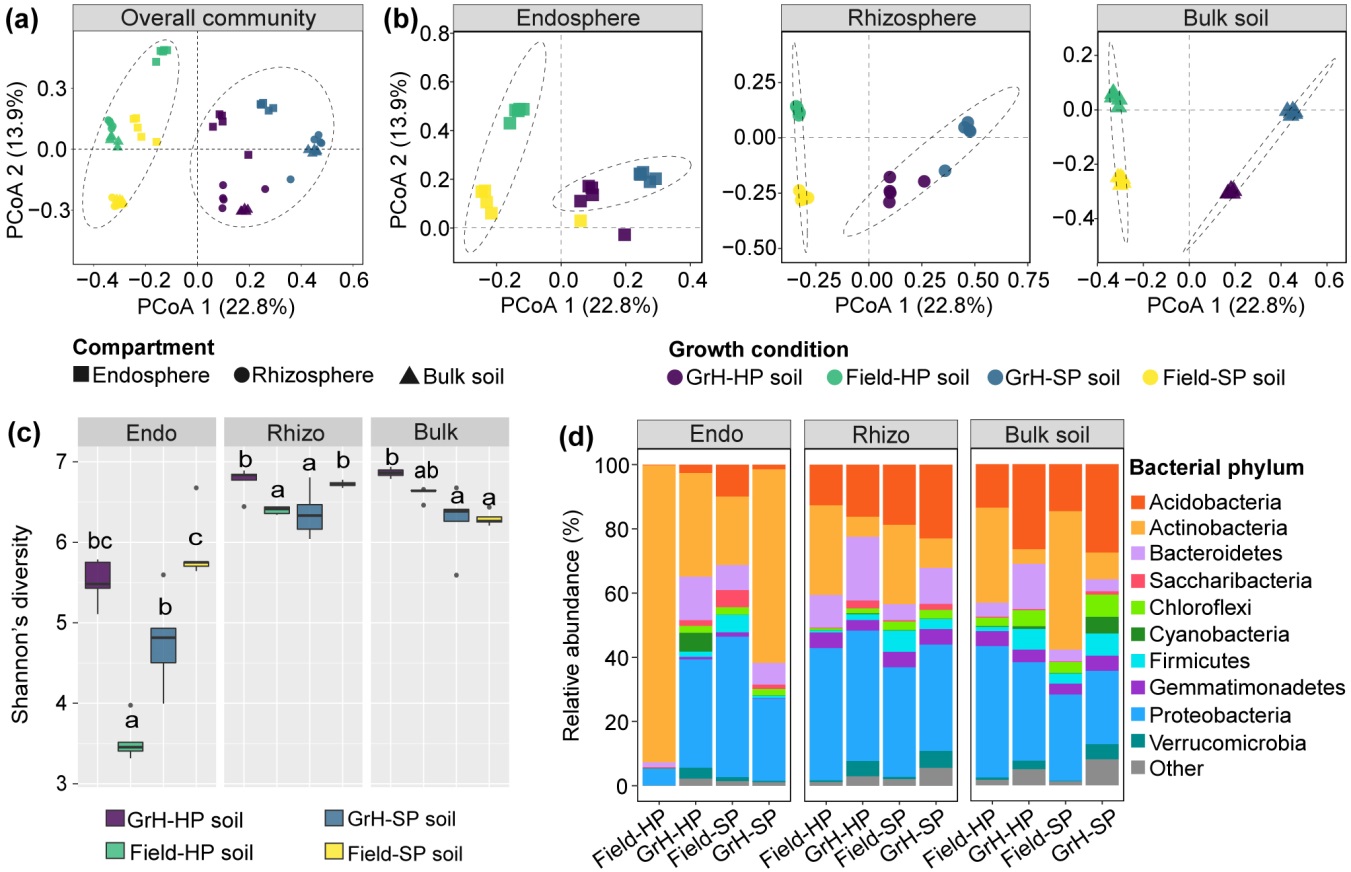

**Figure S2** Structure, diversity and relative abundance of root-associated bacterial communities established under controlled greenhouse and field conditions. PCoA plot using Bray-Curtis dissimilarity depicts the impact of plant growing environment on the structure of the overall bacterial communities (**a**), and bacterial communities inhabiting the bulk soil, rhizosphere and root endosphere of *S. alfredii* grown in HP and SP soils (**b**). Bacterial α-diversity estimated by Shannon’s diversity within each compartment varied by plant growing environment (**c**). The relative abundance of the ten most abundant bacterial phyla within distinct root compartments differed under greenhouse and field conditions. Abbreviation: Endo, root endosphere; Rhizo, rhizosphere; GrH, greenhouse

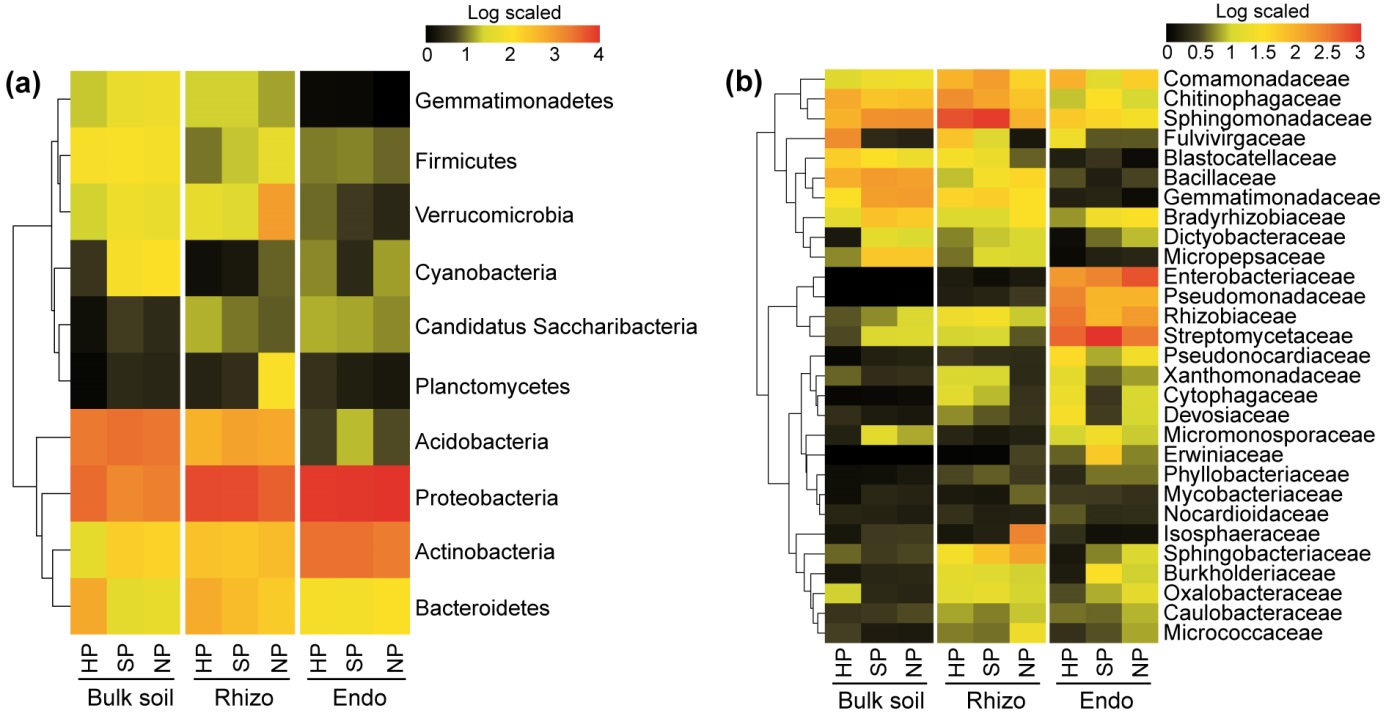

**Figure S3** Plant growth in different soil types under greenhouse conditions affects bacterial community composition in the bulk soil, rhizosphere and root endosphere. (a) Phylum and (b) family distribution of abundant bacterial taxa detected in distinct compartment samples across different soils. Abbreviations: HP, highly polluted soil; SP, slightly polluted soil. NP, non-polluted soil; Rhizo, rhizosphere; Endo, root endosphere.

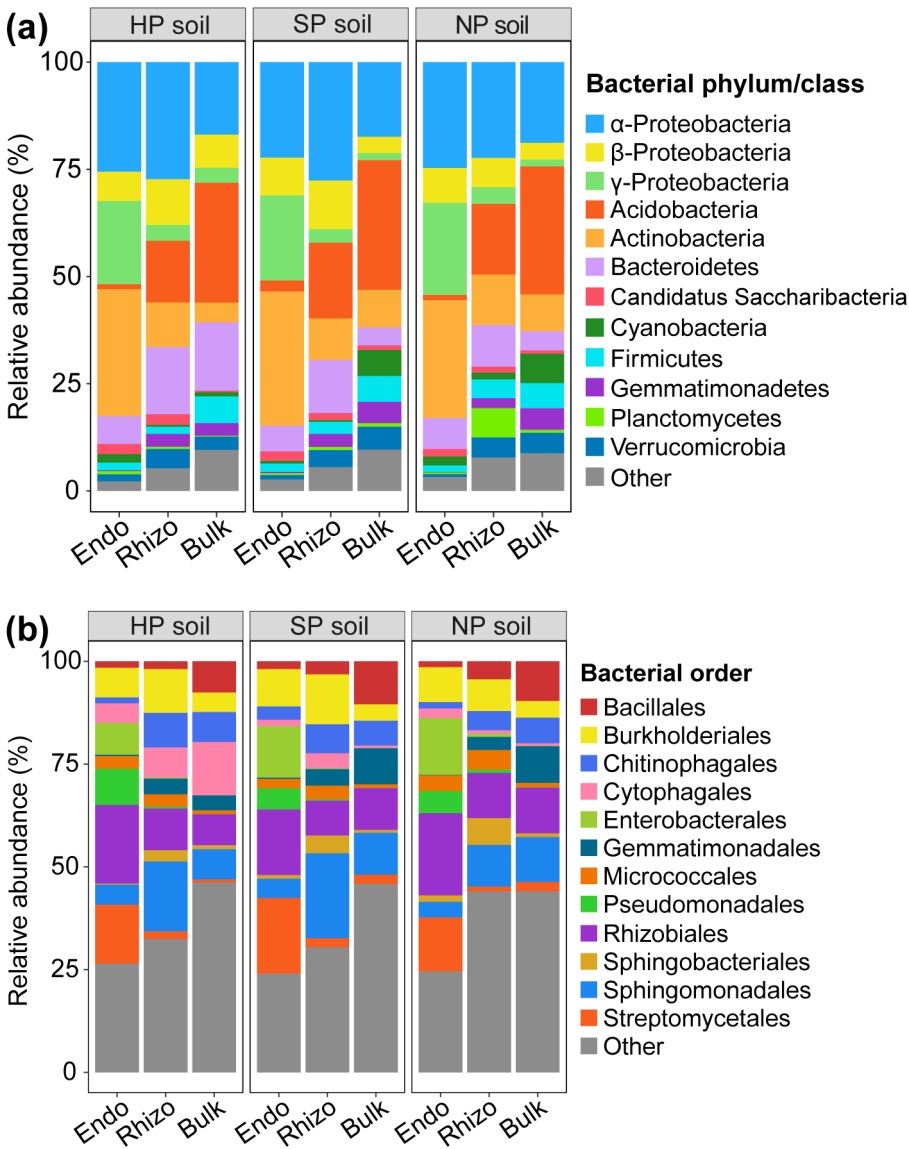

**Figure S4** Distributions of bacterial phyla (**a**) and orders (**b**) inhabiting the bulk soil, rhizosphere and root endosphere across the three distinct soils.

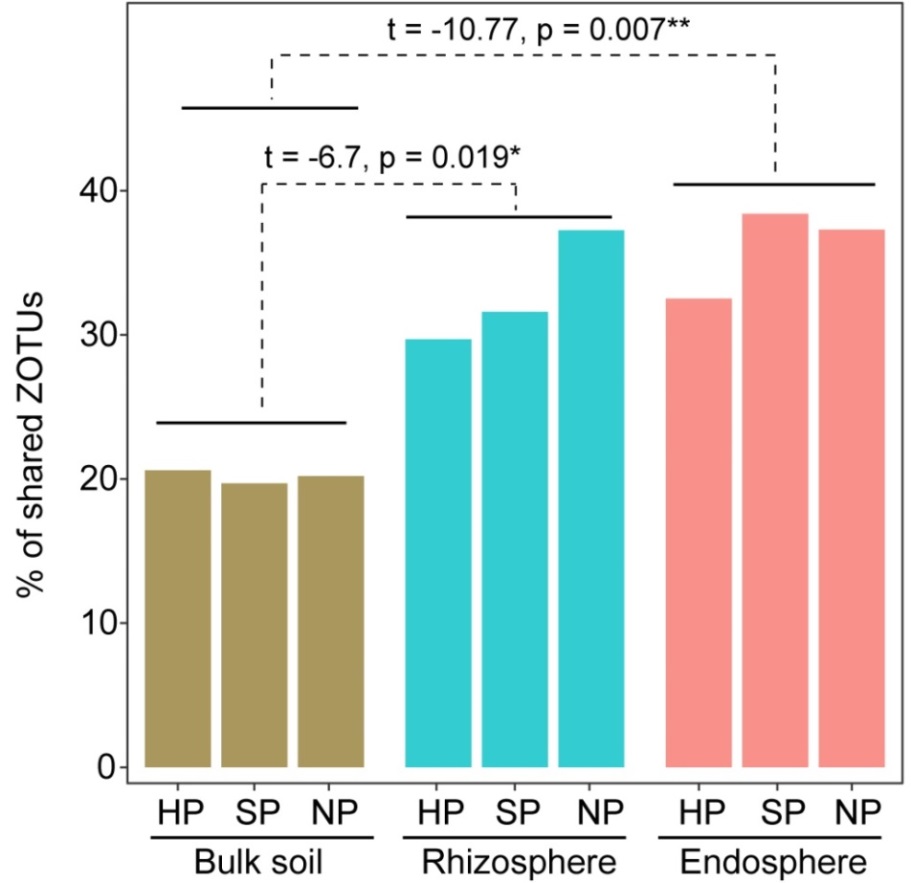

**Figure S5** Percentage of shared abundant ZOTUs (> 20 sequences) among replicate samples in each soil across the root endosphere, rhizosphere and bulk soil. Only ZOTUs containing more than 20 sequence reads were considered. Statistical significance was tested using an independent *t*-test.

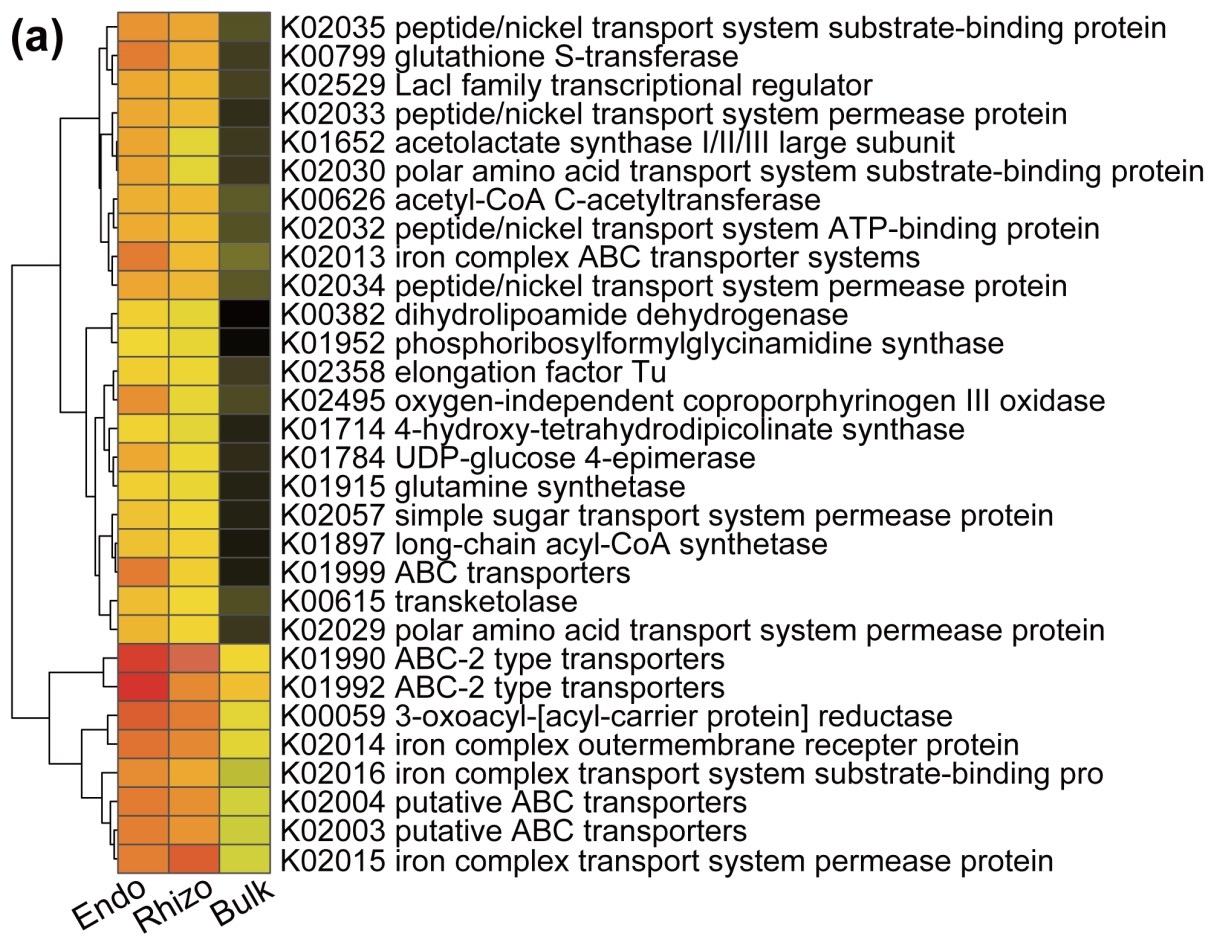

**Figure S6** Heatmap showing the mean relative abundance of the 30 most abundant predicted KEGG functional orthologs of generalist microbiota inhabiting different root compartments.

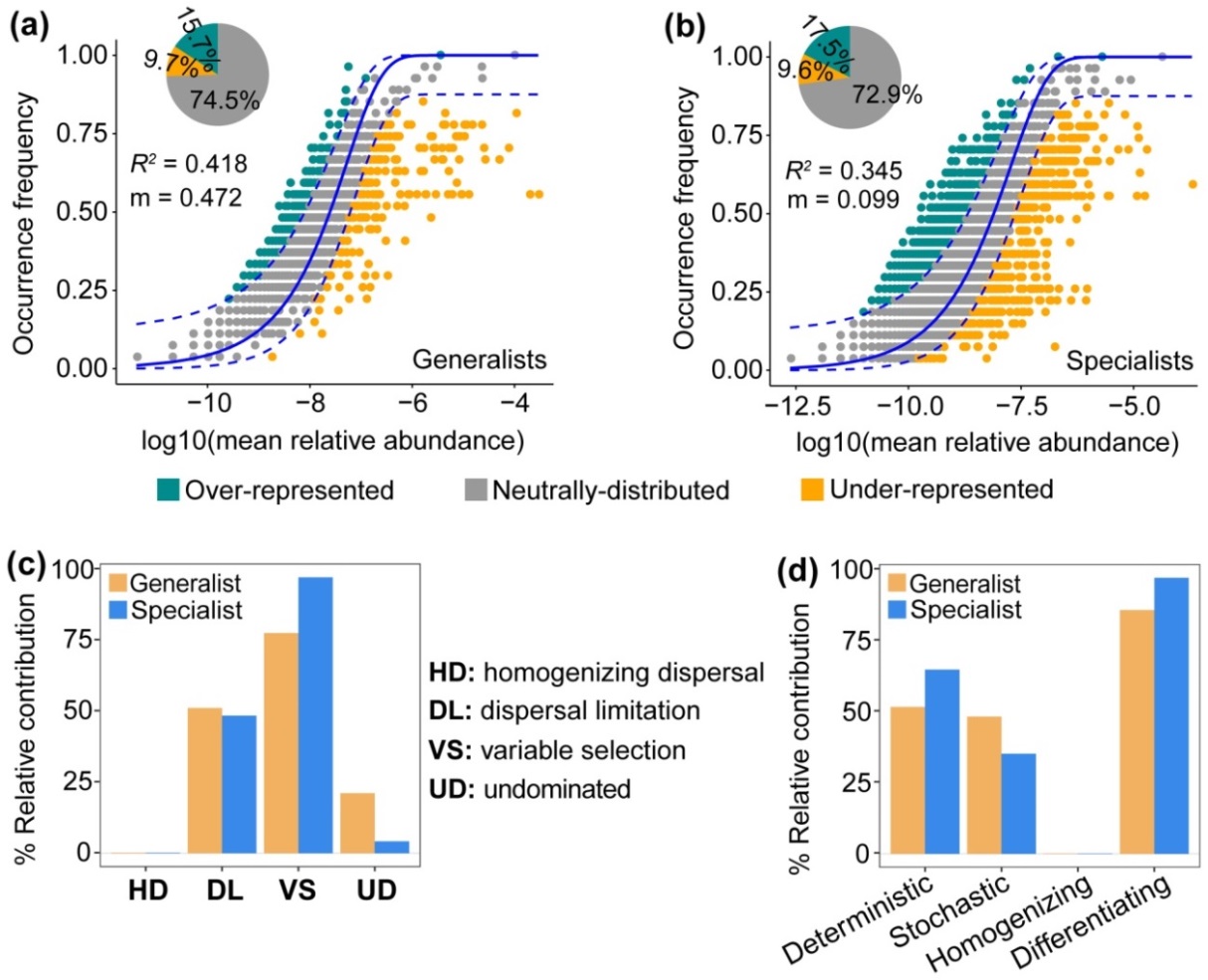

**Figure S7** Community assembly patterns of bacterial sub-communities in the bulk soil under greenhouse conditions. Neutral model applied to assess the effects of random dispersal and ecological drift on the assembly of generalists (**a**) and specialists (**b**) in bulk soil. The solid blue lines indicate the best-fit to the neutral model, and dashed blue lines represent 95% confidence intervals around the model prediction. The pie charts depict the ratio of the over-represented, neutrally distributed and under-represented OTUs in habitat generalists–specialists. (**c**) Null model applied to assess the effects of variable selection, dispersal limitation, homogenizing dispersal and undominated of the bulk soil generalists and specialists. (**d**) The relative contribution of deterministic, stochastic, homogenizing and differentiating processes of the bulk soil generalists and specialists. Deterministic = Variable selection; Stochastic = Dispersal limitation + Homogenizing dispersal; Homogenizing = Homogenizing dispersal; Differentiating = Variable selection + Dispersal limitation.

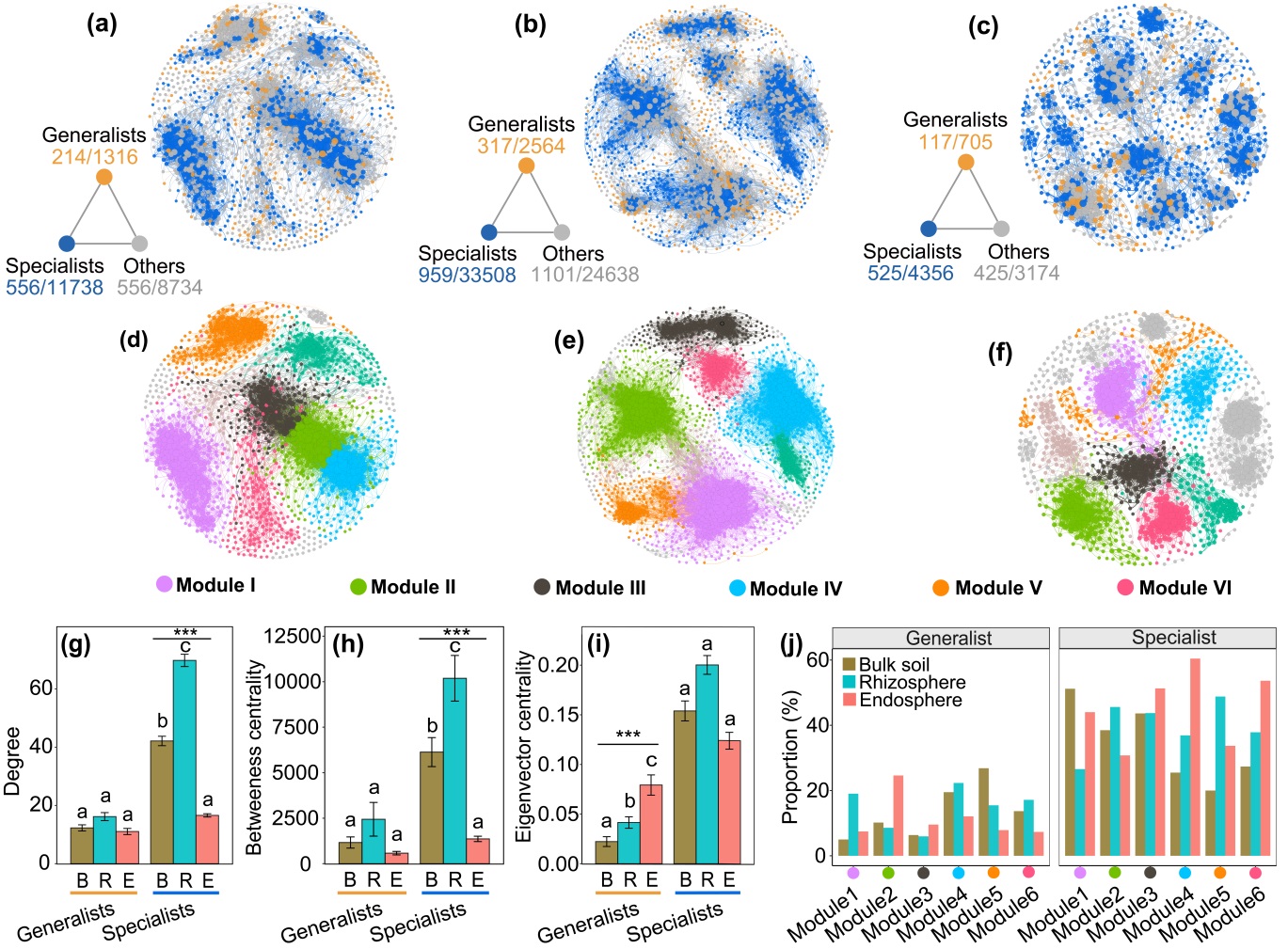

**Figure S8** Co-occurrence patterns of the habitat generalists and specialists across different root compartments. Co-occurrence networks of bacterial taxa in bulk soil (**a**), rhizosphere (**b**) and root endosphere (**c**). The triangle figure on the bottom left of each co-occurrence network is a summary of node-edge statistics. Coloured numbers represent the number of nodes and edges belonging to the corresponding category (i.e. there were 214 nodes and 1316 edges for habitat generalists). The coloured co-occurrence networks are shown for habitat generalists and specialists. Others corresponding to the non-significant ZOTUs shown in Figure 2a. The co-occurrence networks among ZOTUs in bulk soil (**d**), rhizosphere (**e**) and root endosphere (**f**) revealed by modularity class analysis. A connection stands for a strong (Spearman’s *ρ* > 0.75) and significant (*p* < 0.01) correlation. The size of each node is proportional to the number of connections (that is, degree). Node-level topological features of the habitat generalists and specialists across root compartments (**g**). Asterisks denote the significance of compartment type according to Kruskal-wallis test (****P* < 0.001), and different letters above bars indicate significant differences by Kruskal-wallis with Dunn test at *P* < 0.05 level. Bar plots showing the ZOTUs proportion of different sub-communities in each of modules (**h**).

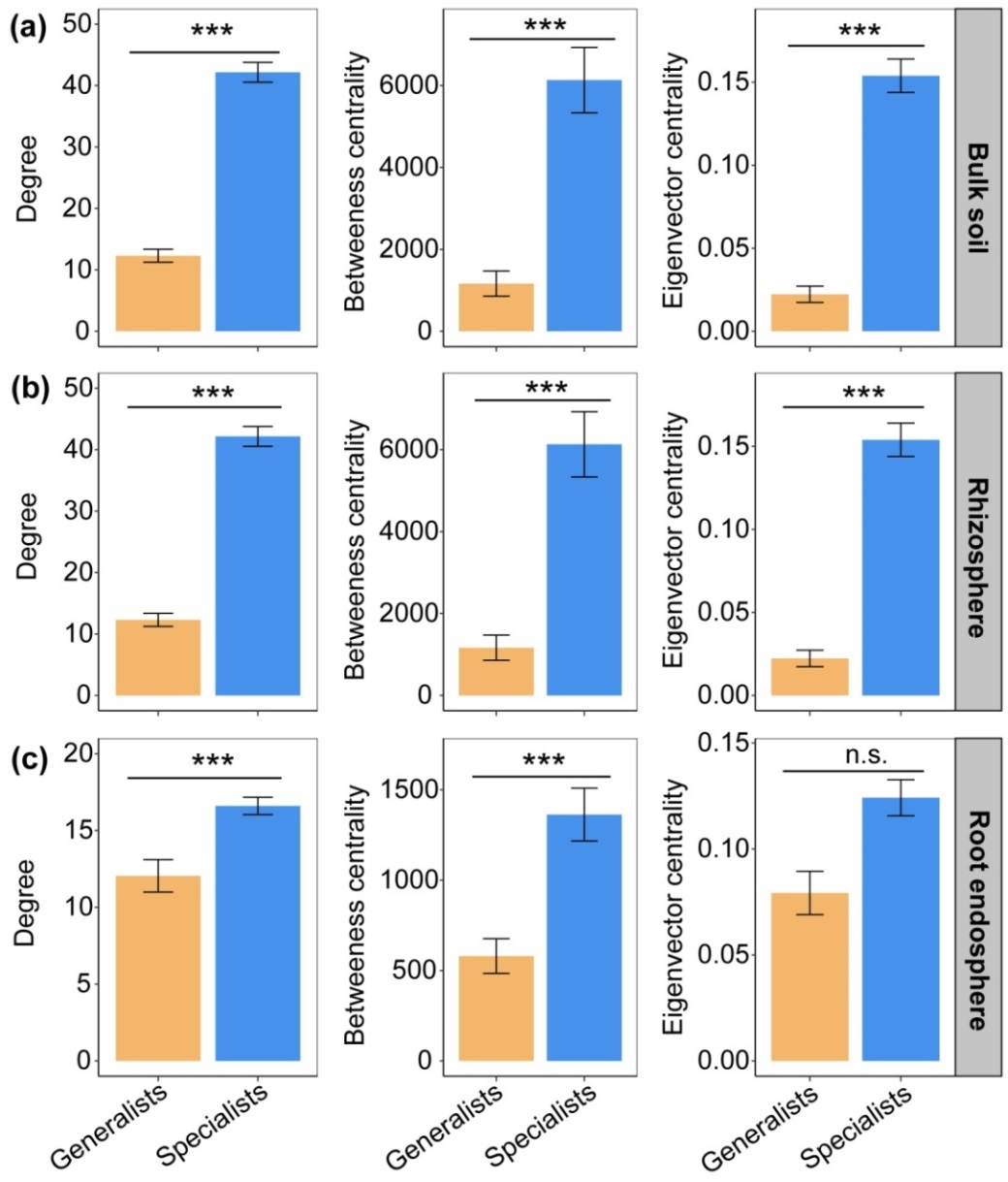

**Figure S9** Node-level topological parameters between the habitat generalists and specialists inhabiting the bulk soil (**a**), the rhizosphere (**b**) and the root endosphere (**c**). Asterisks denote significance in Wilcoxon test, ***P* < 0.01, ****P* < 0.001.

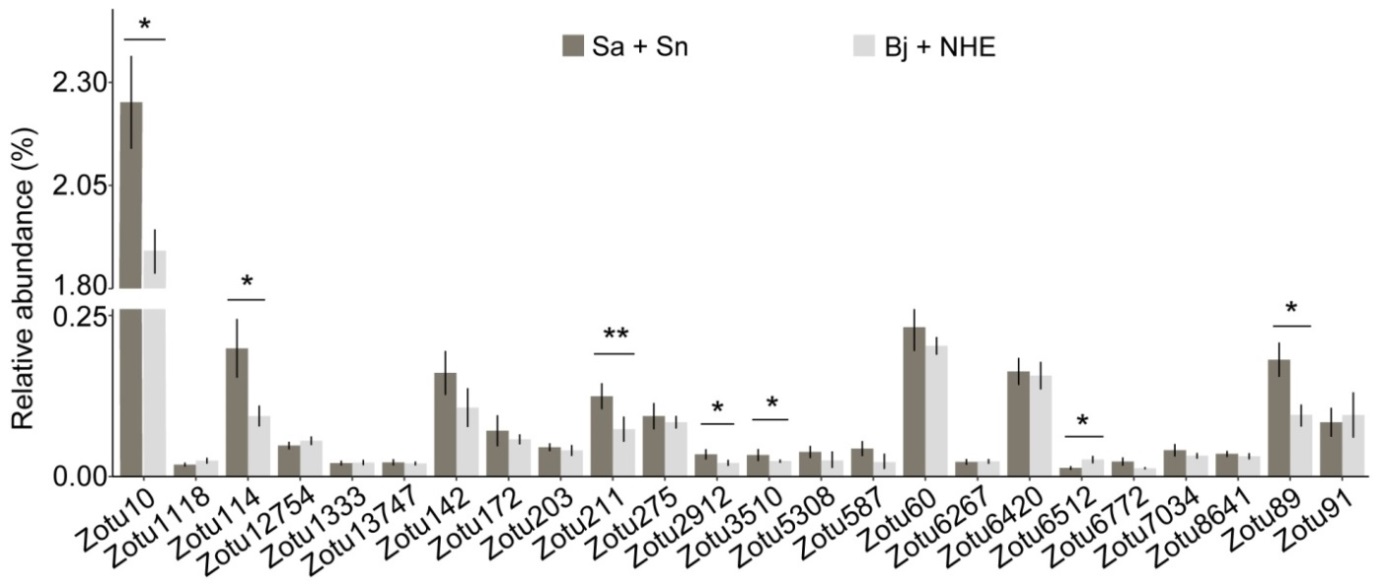

**Figure S10** Comparison of the relative abundance of the 24 generalist core rhizosphere ZOTUs associated with the two plants with stronger Cd/Zn accumulation ability (*S. alfredii* and *S. nigrum*) with the two plants with weaker Cd/Zn accumulation ability (*B. juncea* and non-hyperaccumulator *S. alfredii*) using Wilcoxon test. * *P* < 0.05, ** *P* < 0.01, *** *P* < 0.001. Abbreviation: Sa, *S. alfredii*; Sn, *S. nigrum*; Bj, *B. juncea*; NHE, non-hyperaccumulator *S. alfredii*.

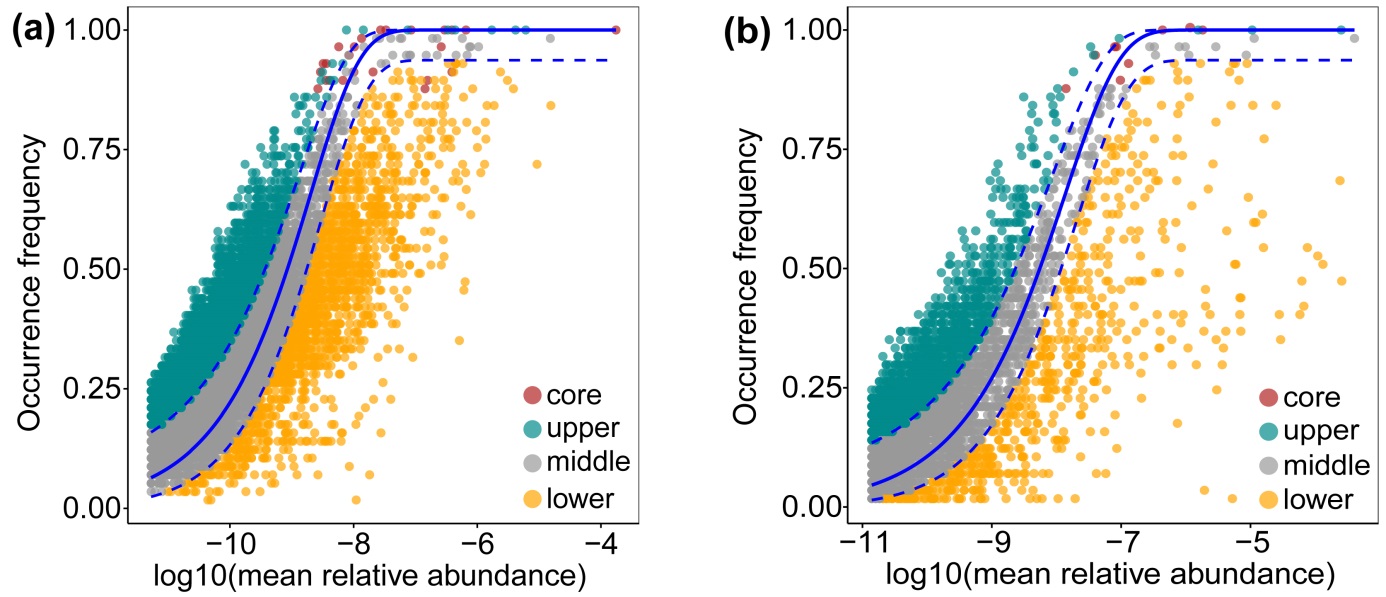

**Figure S11** Neutral model applied to assess the distribution of the generalist core ZOTUs in the rhizosphere (a) and endosphere (b). The solid blue lines indicate the best-fit to the neutral model, and dashed blue lines represent 95% confidence intervals around the model prediction.

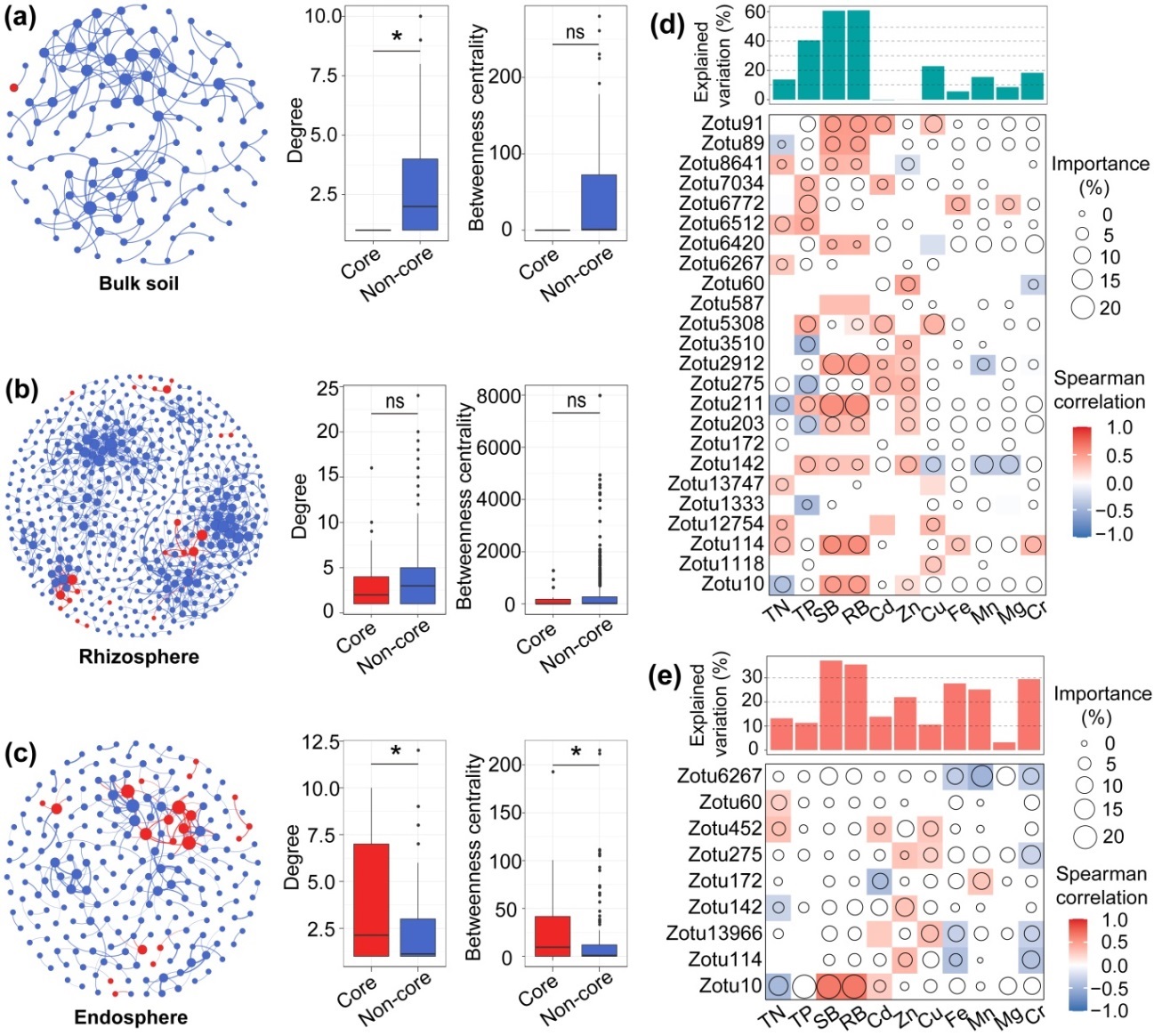

**Figure S12** Bacterial co-occurrence networks of generalist microbiota inhabiting different root compartments and linkages between generalist core microbes and plant functional traits. Co-association networks of generalist core (red) and non-core (blue) taxa and their node-level topological parameters within bulk soil (**a**), rhizosphere (**b**) and root endosphere (**c**). Nodes represent ZOTUs, and edges indicate significant correlation between nodes. The differences in degree and betweenness centrality between generalist core and non-core microbes are based on Wilcoxon test (* *P* < 0.05). Associations between plant functional traits (including shoot and root biomass, plant shoot nutrient and metal accumulation) and the relative abundance of generalist core microbes inhabiting the rhizosphere (d) and root endosphere (e) based on correlation and random forest model. Colours represent Spearman correlations, only significant correlations (*P* < 0.05) are shown. Circle size represents the importance of generalist core microbe (percentage of increased mean square error calculated via random forest model). Abbreviations: SB, shoot biomass; RB, root biomass; TN, total nitrogen; TP, total phosphorus.

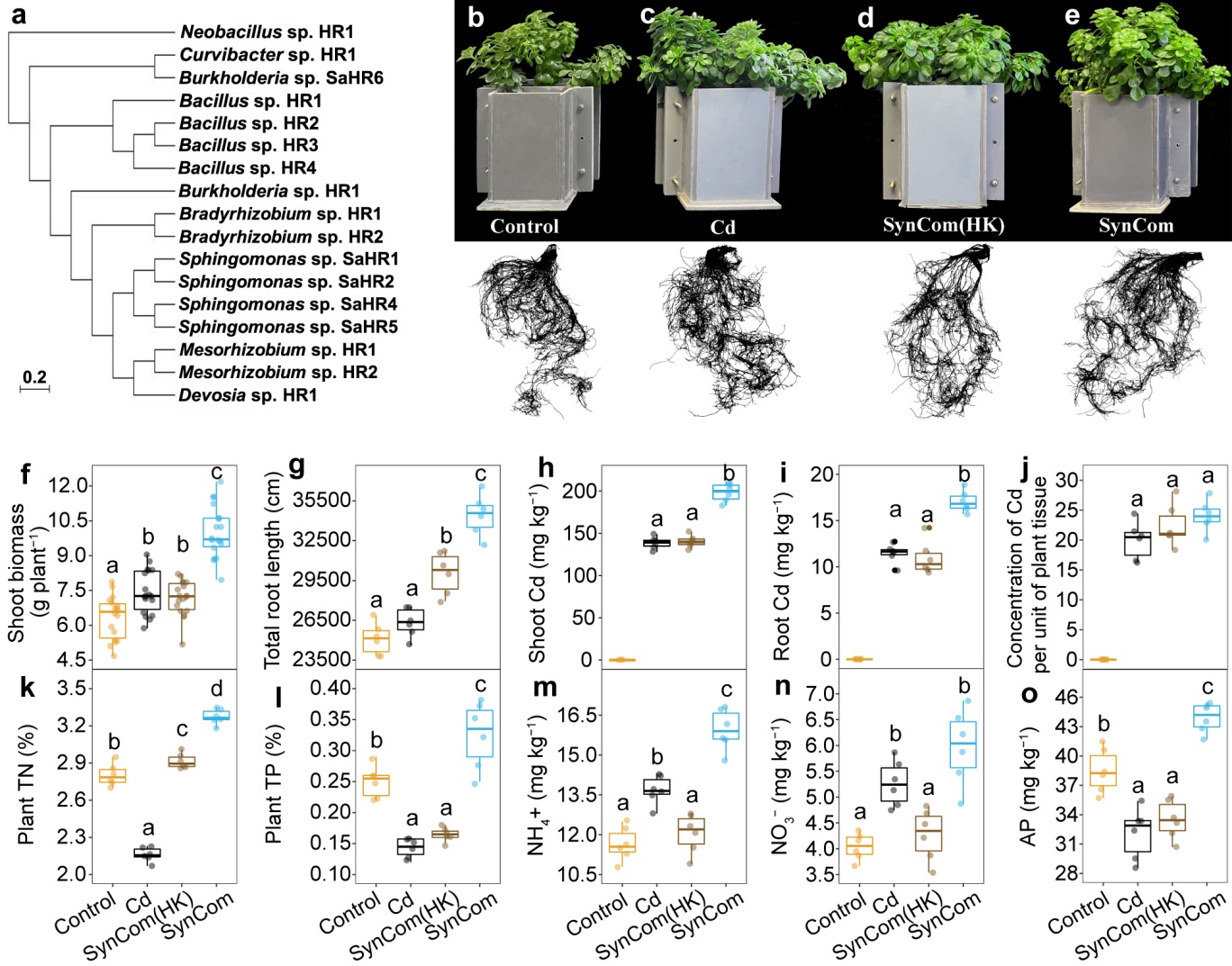

**Figure S13** SynCom comprising the isolates of core rhizospheric bacteria promotes *S. alfredii* growth and improves plant nutrient status. Phylogenetic tree of 17 rhizospheric bacterial isolates used to establish the SynCom, based on the distance values calculated from average nucleotide identity of their full-length 16S rRNA gene with the maximum likelihood method (**a**). Growth status of *S. alfredii* cultivated in different treatments (**b-e**). Effects of Cd, SynCom, heat-killed SynCom [SynCom(HK)] and control (no Cd and SynCom added) treatments on shoot dry weight (**f**), total root length (**g**), concentrations of Cd in shoot (**h**) and root (**i**) and per unit of plant tissue (**j**), plant total nitrogen (**k**) and total phosphorus (**l**), and ammonium (**m**), nitrate (**n**) and available phosphorus (**o**). Values are mean ± s.d. (shown as error bars). Different letters above the box plots indicate significant differences according to Tukey’s HSD test at *P* < 0.05.

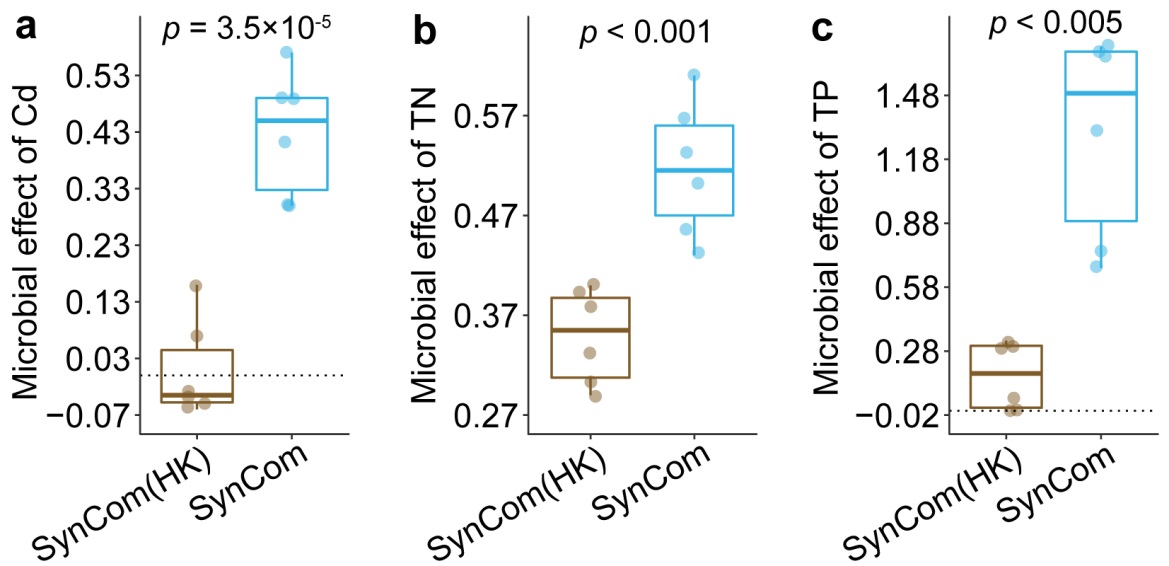

**Figure S14** Microbial effect of rhizospheric SynCom on plant accumulation of Cd, total nitrogen (TN) and total phosphorus (TN) compared with the heat killed SynCom treatment. Pairwise comparison was performed using independent *t* test.

**Tables**

**Table S1** Physiochemical parameters of the three tested soils used in this study

| Soil parameters | Highly polluted soil  (HP soil) | Slightly polluted soil  (SP soil) | Non-polluted soils  (NP soil) |
| --- | --- | --- | --- |
| pH (soil:water=1:2.5) | 6.43 | 6.57 | 4.69 |
| Organic matter (g kg^-1^) | 13.5 | 9.54 | 10.24 |
| NH_4_^+^-N (mg kg^-1^) | 14.8 | 13.9 | 18.7 |
| Total N (g kg^-1^) | 0.75 | 0.57 | 0.95 |
| Available P (mg kg^-1^) | 41.66 | 98.45 | 23.5 |
| Total Cd (mg kg^-1^) | 12.54 | 0.98 | ND |
| Total Zn (mg kg^-1^) | 824.5 | 242.7 | 35.62 |
| DTPA-Cd (mg kg^-1^) | 1.86 | 0.26 | ND |
| DTPA-Zn (mg kg^-1^) | 22.57 | 5.75 | 0.15 |

DTPA solution is a mixture solution of 0.005 M DTPA, 0.01 M CaCl_2_ and 0.1 M TEA. Total N, total nitrogen; P, phosphorus; DTPA-Cd, DTPA-extractable Cd; DTPA-Zn, DTPA-extractable Zn; ND, not detected.

**Table S2** Comparison of bacterial community diversity across plant species, compartments and soils

| Factor | Groups compared | Differences between the groups | | | | | |
| --- | --- | --- | --- | --- | --- | --- | --- |
|  |  | Alpha diversity | | Bray-Curtis distance | | Jaccard distance | |
| **Greenhouse experiment** |  | *R^2^* | *P*-value | *R^2^* | *P*-value | *R^2^* | *P*-value |
| Plant species | Bj *vs* Sa *vs* Sn *vs* NHE | **20.3%** | **< 10^-16^** | **11.7%** | **< 0.001** | **12.1%** | **< 0.001** |
| Compartment | Endo *vs* Rhizo *vs* Bulk | **52.5%** | **< 10^-16^** | **21.7%** | **< 0.001** | **25.3%** | **< 0.001** |
| Soil geographical origin | Hangzhou (HP, SP) *vs* Jian (NP) | 0.2% | NS | **5.9%** | **< 0.001** | **7.4%** | **< 0.001** |
| Soil pollution degree | HP *vs* SP | **1.1%** | **< 0.005** | **8.0%** | **< 0.001** | **11.2%** | **< 0.001** |
| Plant species × Compartment | Interactions between factors | **13.1%** | **< 10^-16^** | **5.1%** | **< 0.001** | **4.0%** | **< 0.001** |
| Compartment × Geographical origin | Interactions between factors | 0.3% | NS | **5.7%** | **< 0.001** | **6.3%** | **< 0.001** |
| Plant species × Geographical origin | Interactions between factors | **2.6%** | **< 10^-6^** | **4.4%** | **< 0.001** | **2.2%** | **< 0.05** |
| **Within the root endosphere compartment** |  |  |  |  |  |  |  |
| Plant species | Bj *vs* Sa *vs* Sn *vs* NHE | **69.4%** | **< 10^-15^** | **34.5%** | **< 0.001** | **33.3%** | **< 0.001** |
| Soil geographical origin | Hangzhou (HP, SP) *vs* Jian (NP) | 0.87% | 0.162 | **4.7%** | **< 0.001** | **4.7%** | **< 0.003** |
| Soil pollution degree | HP *vs* SP | 0.87% | 0.579 | **13.9%** | **< 0.001** | **21.0%** | **< 0.001** |
| **Within the rhizosphere compartment** |  |  |  |  |  |  |  |
| Plant species | Bj *vs* Sa *vs* Sn *vs* NHE | **30.34%** | **< 10^-7^** | **19.9%** | **< 0.001** | **19.2%** | **< 0.001** |
| Soil geographical origin | Hangzhou (HP, SP) *vs* Jian (NP) | **6.44%** | **< 0.005** | **17.0%** | **< 0.001** | **21.6%** | **< 0.001** |
| Soil pollution degree | HP *vs* SP | **10.77%** | **< 0.05** | **21.6%** | **< 0.001** | **36.1%** | **< 0.001** |
| **Within the bulk soil compartment** |  |  |  |  |  |  |  |
| Soil geographical origin | Hangzhou (HP, SP) *vs* Jian (NP) | 7.42% | 0.137 | **36.2%** | **< 0.003** | **53.9%** | **< 0.005** |
| Soil pollution degree | HP *vs* SP | **11.92%** | **< 0.05** | **61.8%** | **< 0.001** | **72.9%** | **< 0.001** |

Abbreviations: Bj, *B. juncea*; Sa, *S. alfredii*; Sn, *S. nigrum*; NHE, non-hyperaccumulating *S. alfredii*; Endo, root endosphere; Rhizo, rhizosphere; HP, highly polluted soil; SP, slightly polluted soil; NP, non-polluted soil.

**Table S3** Comparison of bacterial community diversity across compartments, pollution degree and growing conditions

| Factor | Groups compared | | Differences between the groups | | | |
| --- | --- | --- | --- | --- | --- | --- |
|  |  |  | Alpha diversity | | Bray-Curtis distance | |
| **Field experiment** |  | | *R^2^* | *P*-value | *R^2^* | *P*-value |
| Compartment (C) | Endo vs Rhizo vs Bulk | | **61.6%** | **< 10^-16^** | **10.6%** | **< 0.001** |
| Soil pollution degree (P) | HP vs SP | | 0.3% | NS | **17.0%** | **< 0.001** |
| Plant growing condition (G) | Greenhouse vs Natural | | **0.8%** | **0.021** | **15.3%** | **< 0.001** |
| C × P | Interactions between factors | | **7.5%** | **< 10^-8^** | **14.8%** | **< 0.001** |
| C × G | Interactions between factors | | 0.9% | NS | **7.1%** | **< 0.001** |
| P ×G | Interactions between factors | | **12.7%** | **< 10^-12^** | **8.1%** | **< 0.001** |
| C × P ×G | Interactions between factors | | **10.4%** | **< 10^-10^** | **5.6%** | **< 0.001** |
| **Within the root endosphere compartment** |  | |  |  |  |  |
| Soil pollution degree (P) | HP vs SP | | **17.4%** | **< 0.001** | **18.3%** | **< 0.001** |
| Plant growing condition (G) | Greenhouse vs Natural | | **5.5%** | **0.036** | **23.6%** | **< 0.001** |
| P ×G | Interactions between factors | | **63.7%** | **< 10^-7^** | **19.1%** | **< 0.001** |
| **Within the rhizosphere compartment** |  | |  |  |  |  |
| Soil pollution degree (P) | HP vs SP | | 0.5% | NS | **19.8%** | **< 0.001** |
| Plant growing condition (G) | Greenhouse vs Natural | 0% | | NS | **38.7%** | **< 0.001** |
| P ×G | Interactions between factors | | **56.4%** | **< 0.001** | **22.2%** | **< 0.001** |
| **Within the bulk soil compartment** |  | |  |  |  |  |
| Soil pollution degree (P) | HP vs SP | | **53.9%** | **< 0.001** | **21.2%** | **< 0.001** |
| Plant growing condition (G) | Greenhouse vs Natural | | 3.3% | NS | **40.8%** | **< 0.001** |
| P ×G | Interactions between factors | | 5.3% | NS | **22.2** | **< 0.001** |

**Table S4** The influence of compartment type and soil variation on the relative abundance of the ten most abundant bacterial phyla. (a) Significant differences across compartments and soils (**ANOVA**: aov(Phylum ~ Compartment * Soil variation)). (b) Significant differences between tested soils across compartments (Dunn test). n.s.: non-significant: * *P* < 0.05, ** *P* < 0.01, *** *P* < 0.001 (BH adjusted). HP: highly polluted soil; SP, slightly polluted soil; NP: non-polluted paddy soil.

**(a)**

|  | Compartment | Soil variation | C * S |
| --- | --- | --- | --- |
| Proteobacteria | *** | n.s. | n.s. |
| Actinobacteria | *** | n.s. | n.s. |
| Bacteroidetes | *** | ** | ** |
| Firmicutes | *** | * | * |
| Acidobacteria | *** | n.s. | n.s. |
| Verrucomicrobia | *** | n.s. | n.s. |
| Planctomycetes | *** | *** | *** |
| Gemmatimonadetes | *** | n.s. | * |
| Cyanobacteria | * | n.s. | n.s. |
| Candidatus_Saccharibacteria | *** | ** | ** |

**(b)**

|  | **Bulk soil** | | | **Rhizosphere** | | | **Endosphere** | | |
| --- | --- | --- | --- | --- | --- | --- | --- | --- | --- |
|  | HP-SP | HP-NP | SP-NP | HP-SP | HP-NP | SP-NP | HP-SP | HP-NP | SP-NP |
| Proteobacteria | *** | ** | * | n.s. | * | * | n.s. | n.s. | n.s. |
| Actinobacteria | * | * | n.s. | n.s. | n.s. | n.s. | n.s. | n.s. | n.s. |
| Bacteroidetes | *** | *** | n.s. | n.s. | * | n.s. | n.s. | n.s. | n.s. |
| Firmicutes | n.s. | n.s. | n.s. | n.s. | *** | ** | n.s. | n.s. | n.s. |
| Acidobacteria | n.s. | * | n.s. | n.s. | n.s. | n.s. | n.s. | n.s. | n.s. |
| Verrucomicrobia | ** | * | n.s. | n.s. | n.s. | n.s. | n.s. | * | n.s. |
| Planctomycetes | ** | * | n.s. | n.s. | *** | *** | n.s. | n.s. | n.s. |
| Gemmatimonadetes | * | * | n.s. | n.s. | n.s. | n.s. | n.s. | n.s. | n.s. |
| Cyanobacteria | n.s. | n.s. | * | n.s. | *** | *** | n.s. | n.s. | n.s. |
| Candidatus Saccharibacteria | *** | * | * | n.s. | * | n.s. | n.s. | n.s. | n.s. |

**Table S5** Topological property of co-occurrence networks within different root-associated compartments

| Community | Empirical networks | | | | | | |  | Random networks | | |
| --- | --- | --- | --- | --- | --- | --- | --- | --- | --- | --- | --- |
|  | No. of nodes | No. of edges | Avg. degree | Avg. clustering coefficient | Avg. path length | Diameter | Modularity |  | Avg. clustering coefficient | Avg. path length | Modularity |
| **Endosphere** | 1067 | 8235 | 12.725 | 0.624 | 5.881 | 15 | 0.855 |  | 0.015 | 2.82 | 0.216 |
| **Rhizosphere** | 2378 | 60710 | 41.559 | 0.609 | 6.774 | 17 | 0.676 |  | 0.022 | 2.31 | 0.112 |
| **Bulk soil** | 1636 | 21788 | 28.232 | 0.564 | 9.17 | 25 | 0.664 |  | 0.016 | 2.62 | 0.158 |

**Table S6** Differential abundance analysis of the generalist core ZOTUs between root compartments and bulk soil. Fold change (FC) was determined from the contrasts of the ratio of relative abundance (RA) in root samples to that in bulk soil with the DESeq2 package. lfcSE, standard deviation of log2 FC. *p*-adj, the *p*-values adjusted using the false discovery rate. Only the *p*-adj had a value lower than 0.05 coupled with a log2FC > 0 was shown in bold. The positive value of log2FC means a higher ZOTU RA in rhizosphere and endosphere than in the bulk soil.

| Comparison | | log2FC | | lfcSE | | *p*-value | | *p*-adj |
| --- | --- | --- | --- | --- | --- | --- | --- | --- |
| **Rhizosphere vs. Bulk soil** | | | | | | | | |
| ZOTU10 | | 1.40 | | 0.38 | | 0.0002 | | **0.003** |
| ZOTU1118 | | -1.22 | | 0.38 | | 0.0015 | | 0.013 |
| ZOTU114 | | -0.53 | | 0.36 | | 0.145 | | 0.320 |
| ZOTU12574 | | -0.26 | | 4.449 | | 0.9536 | | >1.000 |
| ZOTU1333 | | 1.257 | | 0.526 | | 0.0168 | | **0.044** |
| ZOTU13747 | | 3.219 | | 0.6404 | | 5.01E-07 | | **1.62E-05** |
| ZOTU142 | | 3.411 | | 0.494 | | 5.05E-12 | | **4.50E-10** |
| ZOTU172 | | 0.307 | | 0.3112 | | 0.3239 | | 0.5138 |
| ZOTU203 | | 1.352 | | 0.378 | | 0.0003 | | **0.0041** |
| ZOTU211 | | -0.5044 | | 0.363 | | 0.16497 | | 0.3460 |
| ZOTU275 | | 1.596 | | 0.3299 | | 1.31E-06 | | **3.84E-05** |
| ZOTU2912 | | 2.606 | | 0.473 | | 3.70E-08 | | **1.59E-06** |
| ZOTU3510 | | 0.857 | | 0.384 | | 0.025 | | 0.0994 |
| ZOTU5308 | | 3.699 | | 0.637 | | 6.21E-09 | | **3.13E-07** |
| ZOTU587 | | -0.825 | | 0.432 | | 0.056 | | 0.1714 |
| ZOTU60 | | -0.836 | | 0.227 | | 0.0002 | | 0.0029 |
| ZOTU6267 | | 3.085 | | 0.663 | | 3.26E-06 | | **8.51E-05** |
| ZOTU6420 | | 0.868 | | 0.321 | | 0.0068 | | **0.039** |
| ZOTU6512 | | 0.765 | | 0.511 | | 0.167 | | 0.349 |
| ZOTU6772 | | 3.645 | | 0.703 | | 2.16E-07 | | **7.89E-06** |
| ZOTU8641 | | -0.833 | | 0.342 | | 0.015 | | 0.068 |
| ZOTU89 | | -1.011 | | 0.543 | | 0.065 | | 0.185 |
| ZOTU91 | | -3.138 | | 0.421 | | 9.35E-14 | | 9.17E-12 |
| ZOTU97 | | 0.656 | | 0.434 | | 0.131 | | 0.300 |
| Comparison | log2FC | | lfcSE | | *p*-value | | *p*-adj | |
| **Endosphere vs. Bulk soil** | | | | | | | | |
| ZOTU114 | | -1.46 | | 0.36 | | 6.35E-05 | | 0.0004 |
| ZOTU275 | | 1.61 | | 0.33 | | 1.25E-06 | | **1.13E-05** |
| ZOTU452 | | 5.67 | | 0.64 | | 6.90E-19 | | **4.16E-17** |
| ZOTU142 | | 2.47 | | 0.50 | | 6.70E-07 | | **6.44E-06** |
| ZOTU60 | | 1.34 | | 0.41 | | 1.15E-05 | | **0.0003** |
| ZOTU13966 | | 4.09 | | 0.63 | | 6.73E-11 | | **1.37E-09** |
| ZOTU172 | | 1.75 | | 0.31 | | 1.51E-08 | | **2.02E-07** |
| ZOTU6267 | | 4.72 | | 0.66 | | 7.95E-13 | | **2.16E-11** |
| ZOTU10 | | -1.66 | | 0.38 | | 1.00E-05 | | 7.47E-05 |

**Table S7** Forward and reverse primers used in quantitative PCR analysis.

| Gene name | Forward primer (5’-3’) | Reverse primer (5’-3’) |
| --- | --- | --- |
| *SaZIP*1 | AGGTG TTATC GGTAT CTGCT | ACTTA TCCCA CGGAG TTTCA |
| *SaZIP*2 | GATTT GACGG AGAAG GAGTA | GTTTG AAGCG TTGCC TGATG |
| *SaZIP*3 | TCCCA CTCGG TCATT ATCGG | TTGGT GTTGT GATGG CGAAA |
| *SaIRT*1 | TGCTC CTGCT TCCGT TCA | TGAAC GGAAG CAGGA GCA |
| *SaNRAMP*3 | AAGAA GCAGC TCATG GGTGT | TAAGC TGCGG TGAAG GTTGA |
| *SaNRAMP*6 | CGTGT GACAT ACCTG AAGTG AT | GCAAG AAGAA CCAAT GTACT GAG |
| *SaHMA*2 | CTCAA AATGC TGCGA AGCGA AG | CTACG CTCTT ACAAC CATGC CTCG |
| *SaHMA*3 | GTTGG CTTTC GCTGG CTATG C | TTCTC AATGT CCGCA ACAGG |
| *ACTIN* | TGTGC TTTCC CTCTA TGCC | CGCTC AGCAG TGGTT GTG |
